# Restoration of metabolic functional metrics from label-free, two-photon cervical tissue images using multiscale deep-learning-based denoising algorithms

**DOI:** 10.1101/2023.06.07.544033

**Authors:** Nilay Vora, Christopher M. Polleys, Filippos Sakellariou, Georgios Georgalis, Hong-Thao Thieu, Elizabeth M. Genega, Narges Jahanseir, Abani Patra, Eric Miller, Irene Georgakoudi

## Abstract

Label-free, two-photon imaging captures morphological and functional metabolic tissue changes and enables enhanced understanding of numerous diseases. However, this modality suffers from low signal arising from limitations imposed by the maximum permissible dose of illumination and the need for rapid image acquisition to avoid motion artifacts. Recently, deep learning methods have been developed to facilitate the extraction of quantitative information from such images. Here, we employ deep neural architectures in the synthesis of a multiscale denoising algorithm optimized for restoring metrics of metabolic activity from low-SNR, two-photon images. Two-photon excited fluorescence (TPEF) images of reduced nicotinamide adenine dinucleotide (phosphate) (NAD(P)H) and flavoproteins (FAD) from freshly excised human cervical tissues are used. We assess the impact of the specific denoising model, loss function, data transformation, and training dataset on established metrics of image restoration when comparing denoised single frame images with corresponding six frame averages, considered as the ground truth. We further assess the restoration accuracy of six metrics of metabolic function from the denoised images relative to ground truth images. Using a novel algorithm based on deep denoising in the wavelet transform domain, we demonstrate optimal recovery of metabolic function metrics. Our results highlight the promise of denoising algorithms to recover diagnostically useful information from low SNR label-free two-photon images and their potential importance in the clinical translation of such imaging.

## Introduction

Metabolism refers to the set of chemical reactions that occur within a cell to produce energy and to build the necessary macromolecules to sustain life^1^. The energetic and macromolecular demands of a cell often change with aging and the onset of several diseases, including cancer, diabetes, neurodegenerative disorders, and cardiovascular diseases^2^. Therefore, it is clear that understanding the nature of such metabolic changes at the cellular level to characterize heterogeneity and dynamic interactions among different cell populations is critical for the development of improved diagnostic and treatment methods^3^. However, established methods to assess metabolic function in the clinic and the laboratory either lack resolution^4^ or are destructive^5^.

One approach that is capable of probing tissue metabolic state with high three-dimensional resolution in a non-destructive manner is two-photon excited fluorescence (TPEF) microscopy^6^. TPEF is a non-linear imaging technique that benefits from intrinsic optical sectioning and the ability to penetrate hundreds of micrometers into bulk tissue^7^. TPEF is also uniquely suited to capture images from endogenous fluorophores such as NAD(P)H and FAD^8^. NADH and FAD are coenzymes that facilitate energy generation and biomolecular synthesis via a number of pathways^9^. Several of these pathways, including the tricarboxylic acid cycle, glutaminolysis, fatty acid oxidation, and oxidative phosphorylation occur in the mitochondria^10^. NADPH plays an important role in anti-oxidant pathways and has similar fluorescence characteristics to those of NADH^11^. Thus, the term NAD(P)H is used throughout this paper to refer to the fluorescence of both NADH and NADPH. A large fraction of the flavin-associated cellular fluorescence is attributed to FAD bound to lipoamide dehydrogenase (LipDH), even though contributions from free FAD and FAD bound to Complex II (electron transfer flavoprotein) may also be significant. Here, we use the term FAD to refer to all flavin-associated fluorescence detected from cells.

Despite the lack of specificity in the origins of the fluorescence signals, the ratio of FAD/NAD(P)H or its normalized definition of FAD/(NAD(P)H+FAD) have been shown to correlate to the oxido-reductive state of the cells in many studies^12–15^. Mitochondria are also characterized by the ability to fuse and fission to enhance energy production and delivery in response to stress or to facilitate removal of damaged mitochondria^16^. Such differences in mitochondrial organization have also been quantified based on analysis of NAD(P)H TPEF images^17, 18^. NAD(P)H fluoresces more efficiently when bound to enzymes typically in the mitochondria; therefore variations in NAD(P)H TPEF intensity fluctuations can be exploited for label-free quantitative assessments of mitochondrial organization (clustering) in cells, tissues, and living humans^17, 19^. Changes in mitochondrial organization have in turn been attributed to metabolic function changes^20–22^. The heterogeneity of parameters such as the redox ratio and mitochondrial clustering within a tissue have also been identified as important indicators of metabolic state^23–25^. A number of studies have already highlighted the diagnostic potential of such assessments in living humans and there is growing interest in performing such measurements at the bedside or via endoscopes to expand the range of diagnostic applications to several organs beyond the skin^26–29^. Fast image acquisition in these settings is critical; however, endoscope designs typically include relatively low numerical aperture (NA) (0.5-0.7) objectives and are not as efficient in the generation and collection of TPEF^30^. As a result, low resolution, noise, and other degradations may mask the diagnostically useful functional features. Thus, approaches to enhance label-free, TPEF images could play a transformative role in the successful translation of this technique to improve tissue metabolic function assessments in the context of diagnosis or treatment.

Traditionally, both standard image processing methods as well as inverse techniques have been used to enhancing the interpretability of TPEF data^31^. These methods are most appropriate when one can easily model the physics associated with the sensing modality and the stochasticity of the data is captured in a computationally convenient distribution. Neither is the case for TPEF sensing where the interaction of light with tissue leads to a highly complex forward model and the data are a mix of Poisson statistics and additive Gaussian noise^32^. Motivated by these challenges as well as the recent success of machine learning methods for addressing a range of image analysis and interpretation problems, we consider the use of deep-learning methods for enhancing TPEF images to improve the extraction of metabolically-relevant information.

Deep-learning-based methods have already been shown to enhance quality and resolution of a wide range of images, including label-free two-photon images^33–37^. Convolutional neural network-based content-aware image restoration (CARE), residual channel attention networks (RCAN), and super-resolution generative adversarial networks (SRGAN) have been developed for this purpose^33–35^. While these models have been applied to fluorescence microscopy data, their use has been limited to exogenously labeled samples which have enhanced contrast compared to label-free images. However, recently, *Shen et al.* (2022) demonstrated the application of a generative adversarial networks (GAN) for the restoration of label-free multimodal nonlinear images^36^. We note that in these and related studies, standard metrics, such as peak SNR (PSNR) and structural similarity index measure (SSIM) are used widely as indicators of the quality of image restoration, even though they may not always match the human visual system’s assessment of image quality (MOS)^33, 34, 36, 37^.

Here, we report on the ability of deep-learning based denoising approaches to restore functional metabolic metrics extracted from label-free TPEF images. Specifically, we consider recovery of average and depth dependent variations in the redox ratio (FAD/(NAD(P)H+FAD)) and mitochondrial clustering extracted from analysis of TPEF images acquired from freshly excised human cervical epithelia, including healthy and precancerous lesions. In addition, we assess whether PSNR and SSIM improvements are correlated with the restoration of the functional metabolic metrics. We consider CARE (a U-net), GANs (SRGAN), and RCAN networks, and assess five loss functions, including mean average error (MAE), mean square error (MSE), SSIM, frequency focal loss (FFL), coefficient of variation (R2), and three combinations of these loss functions (see Supplementary Discussion S1 and Supplementary Fig. S2 online). We also examine whether training on FAD or NAD(P)H images impacts the successful restoration of metabolic function metrics from the corresponding denoised images.

We find that a novel combination of a one level wavelet transformation and CARE models trained to denoise each of the four wavelet domain sub-bands yields denoised images that enable optimized recovery of all metabolic function metrics. Interestingly, we observe that the architecture most successful in recovering metabolic metrics is not optimal in terms of more standard metrics such as PSNR and SSIM used to measure performance. Thus, our results indicate that deep-learning based denoising algorithms may require distinct multiscale training and testing approaches for the recovery of functional metrics needed for improved diagnosis and for understanding the drivers of disease and development of novel therapeutics, instead of traditional morphological image quality metrics.

## Results

### Identification of the optimal deep-learning model architecture for denoising label-free, optical TPEF images to enable recovery of metabolic function metrics

Human cervical tissue biopsies were collected from 54 patients and imaged immediately upon excision, as described in *Methods: Optical Instrumentation and Image Acquisition* (Figure 1). Several regions of interest (ROIs) were imaged from each biopsy. Multiple optical sections (OS) were imaged from each ROI at distinct depths. At each OS, we acquired TPEF images at a combination of two excitation wavelengths (775 and 860 nm) and three or four emission bands. Images collected at 775 nm excitation 435-485 nm emission were attributed primarily to NAD(P)H, while images at 860 nm excitation 500-550 nm emission were considered to contain signal primarily from FAD and FAD bound to lipoamide dehydrogenase. Six frames were acquired at each wavelength setting. To reduce the contribution of noise, these six frames were averaged together. Metrics extracted from these averaged images were previously observed to enable highly sensitive and specific detection of cervical pre-cancer^25^. The averaged image was therefore considered the ground truth used for training and testing the denoising success of single frames. Single frames, the corresponding denoised, and ground truth images were analyzed using established procedures to extract the redox ratio (RR), defined as FAD/(NAD(P)H+FAD) in this study, and mitochondrial clustering (β) (**Figure 1**). All models (**Figure 1**) were trained and evaluated with identically generated image stacks. Various combinations of model architectures, loss functions, data transformations, and training data combinations, as outlined in Table 1, were evaluated on 3229 total OSs representing healthy/benign cervical tissues as well as precancerous (low-grade and high grade) squamous intra-epithelial lesions (LSIL and HSIL, respectively).

**Figure 1:**
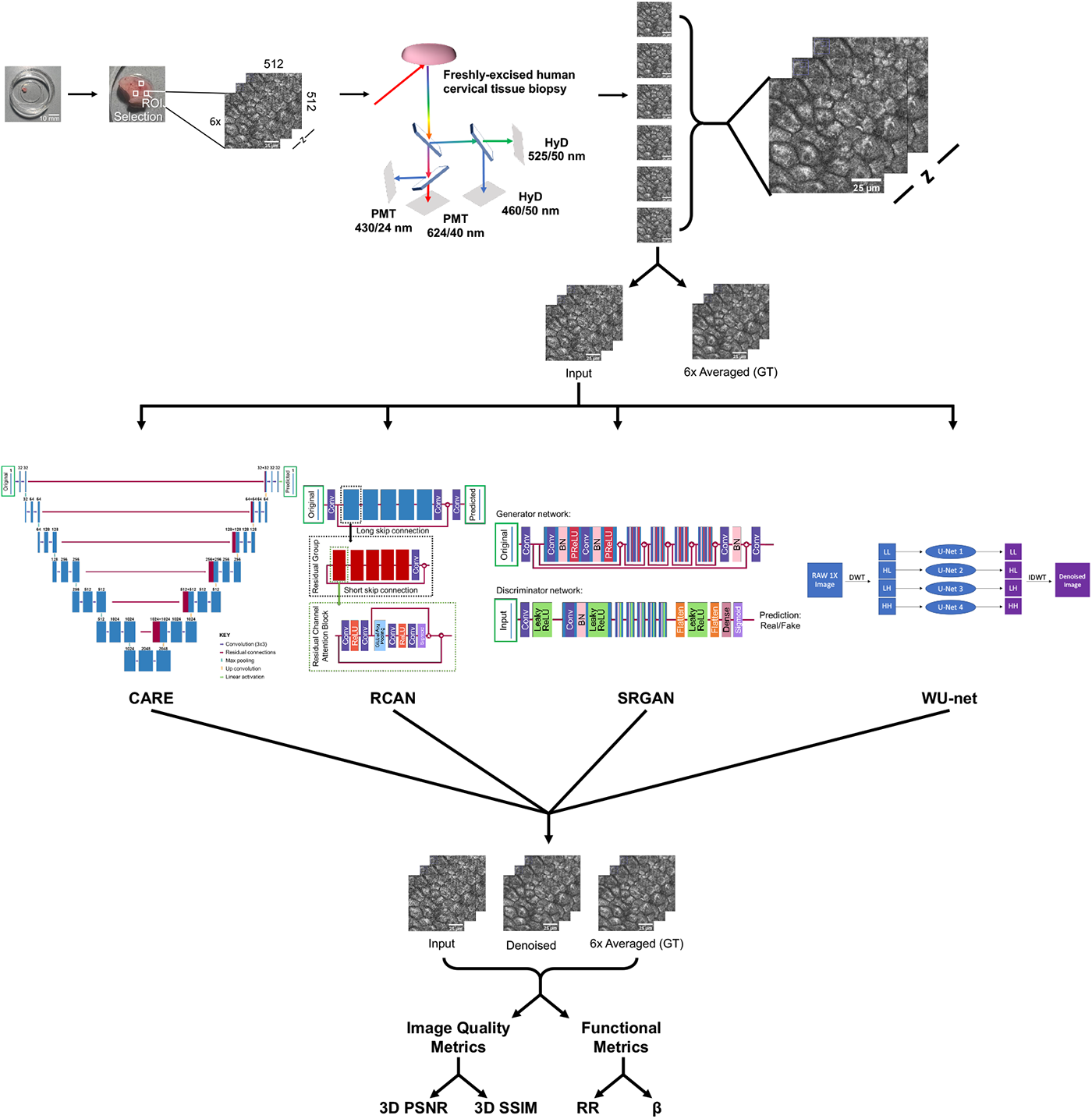
Summary of deep learning pipeline. Human cervical tissue biopsies are collected and subsequently imaged within 4 hours post-excision. Collected biopsies are plated on glass bottom dishes and imaged using a Leica SP8 commercial microscope. At a minimum, three ROIs are imaged per sample. At each ROI, multiple optical sections are imaged at distinct depths through the epithelium. Depth-resolved, two-photon OSs are collected using 2 excitation wavelengths and several bandpass-filtered detectors. Six images are captured for each excitation/emission wavelength and every OS at a given depth, z. These six images are averaged together to generate the ground truth image set. A random image from the six per depth z is selected as the input (RAW) image. The paired image stacks are provided to the neural network for training and denoising. Four-leading denoising networks are used in this study to denoise input images: a previously described CARE model, an RCAN model, an SRGAN model, and a WU-net^33–35, 37^. Denoised images and Input images are compared against 6x Averaged images to determine 3D PSNR and SSIM along with metabolic metrics. Scale bar = 25 μm.

PSNR and SSIM improvements are standard metrics of image visual quality and have been used in other studies focused on denoising biomedical images as an indicator of model success^33, 34, 36, 37^. We aimed to assess whether images restored by models that yield optimized PSNR and SSIM values result in accurate recovery of metabolic metrics (Figure 2). For evaluation of model architecture, loss function, and signal type, only results from models trained on NAD(P)H data from tissues of known benign status were included.

**Table 1:** Summary of all parameters explored during training and optimization of the final model (highlighted in bold). Results shown below are focused on the optimized model, but all combinations were trained and evaluated.

| Model Architecture | CARE | RCAN | SRGAN |  |  |
| --- | --- | --- | --- | --- | --- |
| Loss Functions | MAE / L1 | MSE / L2 | SSIM + L2 | SSIM + FFL | **SSIM + R2** |
| Signal Pre-processing Method | **Wavelet Transform** | None |  |  |  |
| Training Data Format | Healthy Data Only | **Healthy and Diseased (Mix)** |  |  |  |
| Training Data Type | NAD(P)H Data | **FAD Data** |  |  |  |

Leading denoising model architectures were selected for evaluation based on a comprehensive literature search. CARE, RCAN, and SRGAN (Figure 1) models were trained as described in *Methods: Deep Learning Model Description* and *Deep Learning Performance Benchmark*. A representative OS from a HSIL biopsy is shown in Figure 3**a**. Results shown were generated by models trained using an SSIM + Mean Squared Error (MSE or L_2_) loss function. A summary of all parameters used to generate the figures and tables are listed in Table 2.

**Figure 2:**
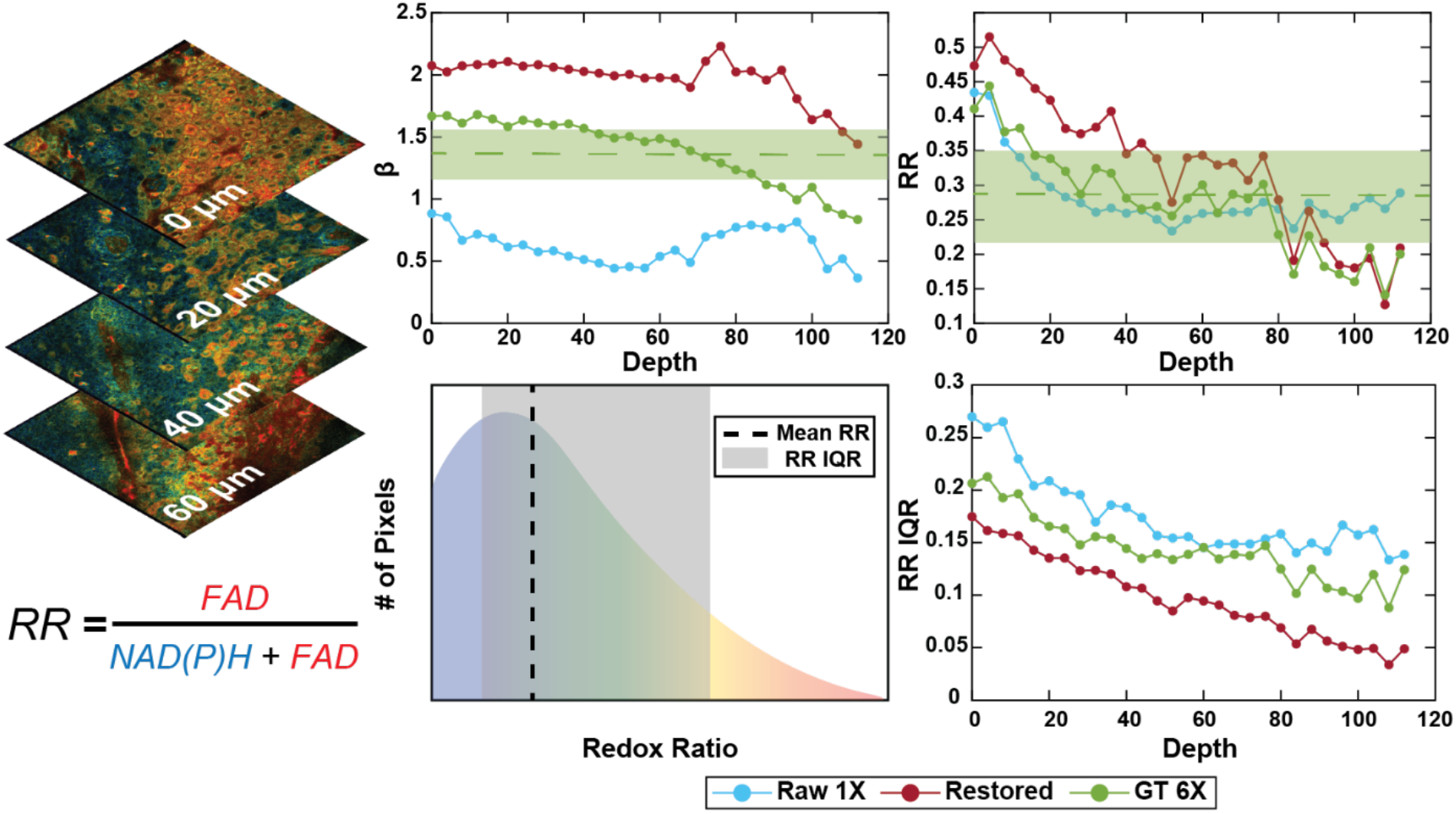
β and RR metrics are extracted from each OS *(See Materials and Methods: Morphological and Functional Metrics* for greater detail). Depth-dependent trends across the multiple cell layers of the cervical squamous epithelium are assessed for input images (RAW 1X), denoised images (Restored), and six-frame averaged, ground truth images (GT 6X). Measurements of mean values and corresponding variability *across all depths* are shown as a dashed line and shaded region in the mitochondrial clustering, β, and RR (FAD/(NAD(P)H+FAD)) panels for the GT 6X image. The distribution of RR values for each OS is used to extract the interquartile range (IQR), representing the range of values within the 25% and 75% of the RR distributions and providing an assessment of intra-field RR heterogeneity. IQR variability is a metric of inter-field (depth-dependent) RR heterogeneity.

Prior to denoising, standard image quality metrics were calculated for input (RAW 1X) images by comparing the RAW 1X images to ground truth (GT 6X) images. PSNR and SSIM values were calculated using the GT 6X image as a reference and RAW 1X or denoised images as the distorted image^38^. Across all images, FAD image PSNR was greater than NAD(P)H image PSNR (Table 3), even though FAD images featured lower cytoplasmic signal compared to NAD(P)H images (Figure 3**a**). During PSNR calculation, the reduced signal intensity led to smaller differences between RAW 1X and GT 6X images and yielded a greater observed PSNR value. This observation was also consistent with results from other studies^36^. SSIM values were consistent between NAD(P)H and FAD images (Table 3). Corruption of the GT 6X images for both channels by noise was expected to have similar effects on structural similarity and calculated SSIM values.

**Table 2:** Summary of parameters used to generate Figures 3-6 and Table 3-6. Parameters are bolded when all combinations from Table 1 are used.

|  | Model Architecture | Loss Functions | Pre-Processing Method | Training Data Format | Training Data Type |
| --- | --- | --- | --- | --- | --- |
| Figure 3 / Table 3 | **All** | SSIM + L2 Loss | None | Healthy Only | NAD(P)H |
| Figure 4 / Table 4 | CARE | SSIM + R2 | **All** | Healthy Only | NAD(P)H |
| Figure 5 / Table 5 | CARE | SSIM + R2 | Wavelet Transform | **All** | **All** |
| Figure 6 / Table 6 | CARE | SSIM + R2<br>MSE<br>MAE | **All** | **All** | **All** |

We used 777 and 109 RAW 1X NAD(P)H OSs for training and validation of the models, respectively. Each 512 x 512 OS was patched into four-256 x 256 image patches (OSP) prior to training and validation (3108 and 436 OSPs, respectively). All three models were trained before being evaluated on an independent set of 2343 OSs (9372 OSPs). Metrics of image quality and metabolic function were calculated as described in *Methods: Deep Learning Metrics* and *Methods: Morphological and Functional Metrics* sections.

**Figure 3:**
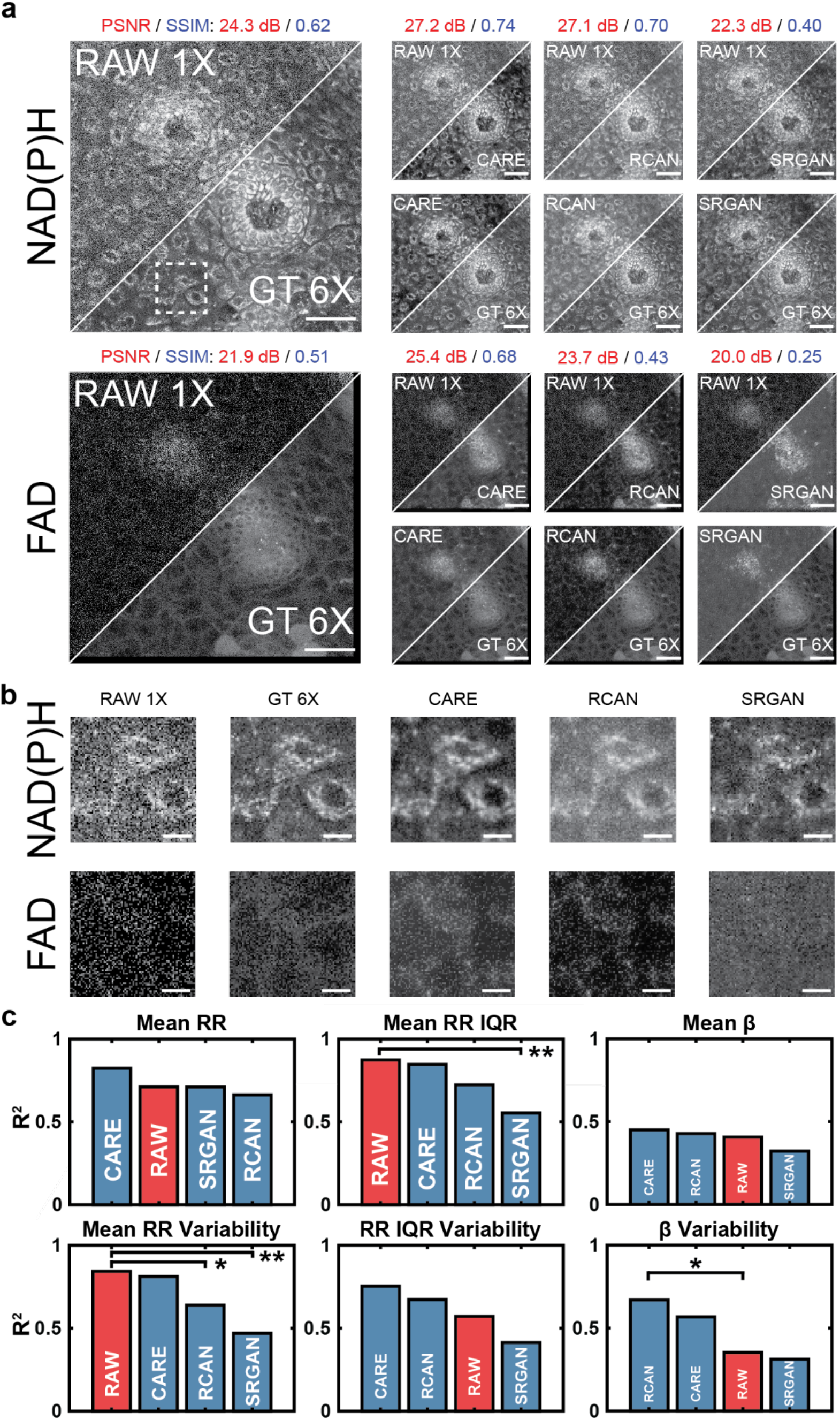
**(a)** A 290 x 290 μm^2^ field of view from a low-grade squamous intraepithelial lesion (LSIL) cervical tissue biopsy. NAD(P)H and FAD images for the same region are shown along with the corresponding denoised image from each of the three trained models (CARE, RCAN, and SRGAN). Scale bar = 50 μm. **(b)** A 44.2 x 44.2 μm^2^ field of view (white square in **a**) of three cells. NAD(P)H and FAD images are shown for all models and the input and ground truth images. Scale bar = 10 μm. **(c)** Bar plots of the coefficient of determination of all downstream metrics for images denoised by all models and RAW 1X vs. the GT 6X image. Fisher r to z transformation was used to measure significance. *p<0.05 and **p<0.01.

CARE-generated image stacks demonstrated higher PSNR for FAD images and higher SSIM for both NAD(P)H and FAD images compared to restored-image stacks generated by RCAN and SRGAN. Across all test set images, standard metrics of image quality (Table 3) and visual inspection (Figure 3**b**) suggested RCAN- and CARE-denoised images had similar image quality. Across the entire test set, we observed SRGAN failed to restore cellular features within the GT 6X images (Figure 3**b**) and underperformed even relative to RAW 1X images in standard image quality metrics (Table 3). Perceptual loss was believed to impact content restoration in the SRGAN architecture^34^. Inputs for perceptual loss calculations have been shown to impact significantly SRGAN performance and were likely the cause of SRGAN’s poor recovery of image quality^34^.

To assess restoration of metabolic activity, depth-dependent optical RR and mitochondrial clustering (β) values were calculated for the restored images, input (RAW 1X) images, and ground truth (GT 6X) images (Figure 2). Pearson correlation coefficient values were calculated between the metabolic function metrics from the GT 6X and either the RAW 1X or restored images. Statistical significance was derived from Fisher-r-to-z transformation for all metrics of interest. Interestingly, analysis of the RAW 1X images led to very high correlations with metrics of RR intra- and inter-field variability compared to GT 6X images. We hypothesized that similar sources of noise in both FAD and NAD(P)H images led to this outcome since RR metrics were calculated using a ratio of FAD and NAD(P)H intensity measurements. It was for this reason that in this initial comparison, we trained models on NAD(P)H images and applied the same weights to NAD(P)H and FAD images. RCAN-generated images demonstrated statistically significant recovery of β variability (*σ*^2^(*B*)) (Figure 3**c**). However, recovery of mean RR variability by this model was poor (Figure 3**c**). CARE-denoised images, overall, featured higher (albeit not statistically significant) correlations with RR metrics compared to all other models (Figure 3**c**). Thus, the U-net architecture of CARE was utilized for all further optimization steps.

**Table 3:**
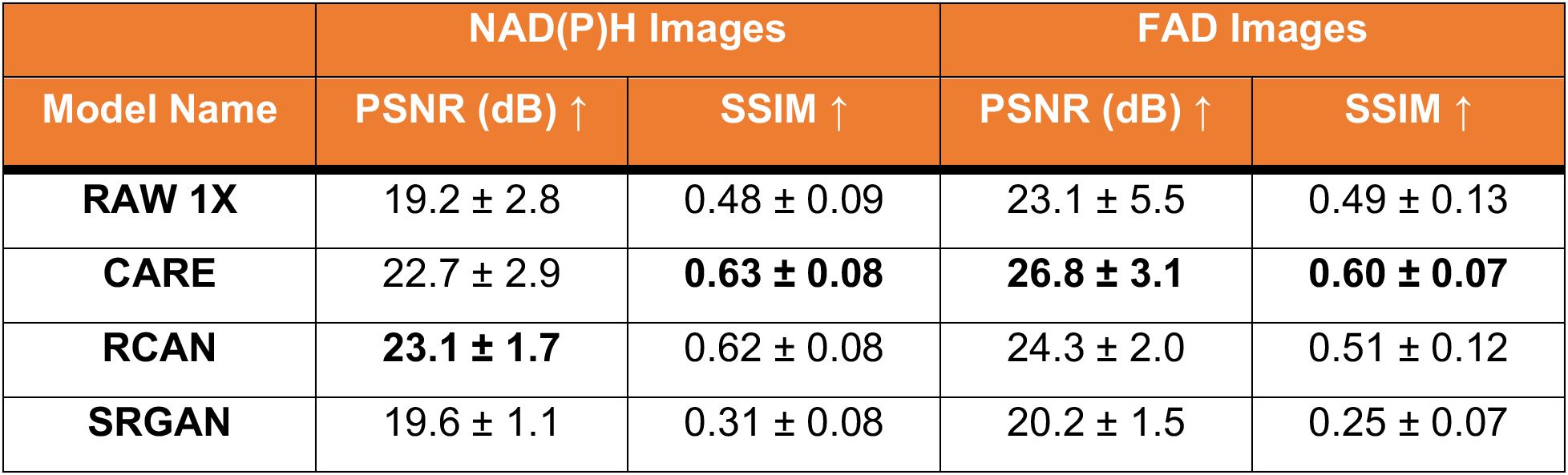
Summary of standard metrics of image quality for RAW 1X images and denoised images generated from various model architectures. Values are reported for mean performance (± standard deviation) across all test set ROIs.

| Model Name | NAD(P)H Images |  | FAD Images |  |
| --- | --- | --- | --- | --- |
| | PSNR (dB) ↑ | SSIM ↑ | PSNR (dB) ↑ | SSIM ↑ |
| **RAW 1X** | 19.2 ± 2.8 | 0.48 ± 0.09 | 23.1 ± 5.5 | 0.49 ± 0.13 |
| **CARE** | 22.7 ± 2.9 | **0.63 ± 0.08** | **26.8 ± 3.1** | **0.60 ± 0.07** |
| **RCAN** | **23.1 ± 1.7** | 0.62 ± 0.08 | 24.3 ± 2.0 | 0.51 ± 0.12 |
| **SRGAN** | 19.6 ± 1.1 | 0.31 ± 0.08 | 20.2 ± 1.5 | 0.25 ± 0.07 |

### A Multiscale Image Transformation Enhances Quantification of Mitochondrial Clustering

Although denoising improved the restoration of *σ*^2^*(β)*, the mean β (β) values of the denoised images were not well correlated with the values from the GT 6X images. We considered discrete wavelet transformation (DWT) to enhance high spatial frequency restoration necessary for β metric calculations. A single level DWT, transformed each image into four sub-band images: a coarser scale approximation and three *detail* images; one horizontal, one vertical, and one diagonal^39^. To generate the three subband images, a basis function, called a mother wavelet, was convolved along both dimensions of the original image while an associated scaling function was used to generate the coarser approximation. During standard wavelet-based denoising, thresholds are used to remove noise from wavelet-transformed detail images, before implementing an inverse-transform to recover the restored image^40, 41^. The DWT has been shown to be advantageous compared to traditional low-pass filtering as the pixel-by-pixel convolution with the mother wavelet preserves correlations of high frequency features. In this study, we used deep learning models trained on each of the transformed images to adaptively learn the best threshold for denoising of low frequencies (approximation) and high frequencies (details) rather than relying on arbitrary thresholding for denoising (see Supplementary Discussion S2 online)^42^. As with any DWT-denoising model, the selection of the correct mother wavelet played a significant role in model performance. For all models, mother wavelets from the biorthogonal, coiflets, and Daubechies families were evaluated. These mother wavelets families were selected due to their frequent use in denoising tasks^43^. Multiple models were trained and evaluated, with biorthogonal 1.1 yielding the highest recovery of metabolic metrics (data not shown). As such, biorthogonal 1.1 was used for all subsequent model optimization.

**Figure 4:**
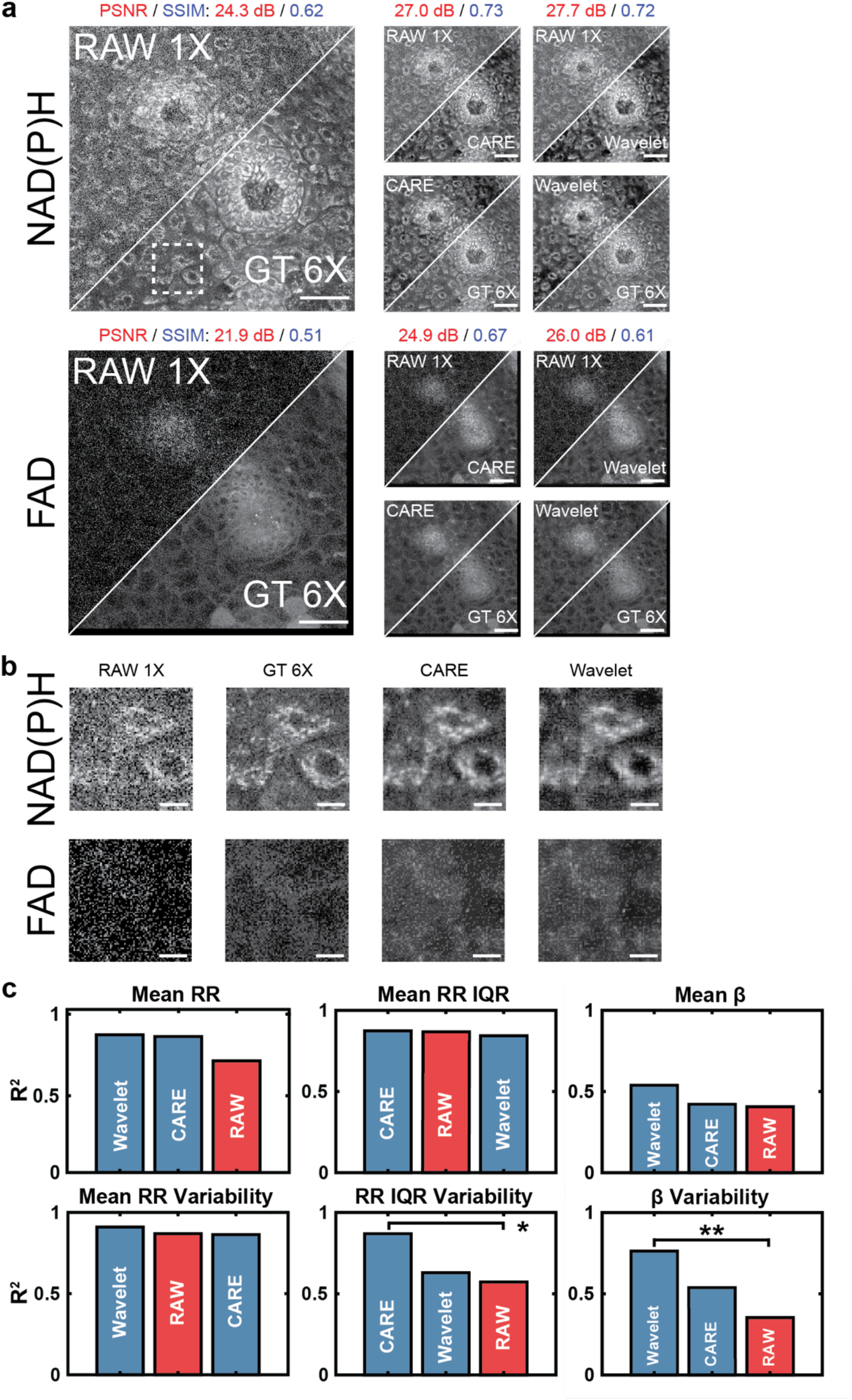
**(a)** A 290 x 290 μm^2^ field of view from a LSIL cervical tissue biopsy. NAD(P)H and FAD images for the same region are shown along with the corresponding denoised image from each signal pre-processing method utilized (Single frame image and Wavelet transformation). Scale bar = 50 μm. **(b)** A 44.2 x 44.2 μm^2^ field of view (white square in **a**) of three cells. NAD(P)H and FAD images are shown for all models based on the corresponding signal pre-processing method used during training and the input and ground truth images. Scale bar = 10 μm. **(c)** Bar plots of the coefficient of determination of all downstream metrics for images denoised by all models trained based on the corresponding signal pre-processing method used during training and RAW 1X vs. the GT 6X image. Fisher r to z transformation was used to measure significance. *p<0.05 and **p<0.01.

Application of DWT before training four CARE models and inverse DWT (iDWT) after evaluation yielded images with improved FAD and NAD(P)H PSNR with slight decreases in SSIM (Figure 4**a**). Across the entire test set, NAD(P)H PSNR improved using WU-net while FAD PSNR and SSIM both decreased compared to CARE (Table 4). All loss functions were evaluated for WU-net, with SSIM + R2 loss (results shown in Figure 4) and SSIM + FFL loss (see Supplementary Table S3 online) yielding the best overall performance. WU-net denoised NAD(P)H images extracted similar cellular structures as the CARE derived images but featured lower background signal and small fluctuations in cytoplasmic signal leading to the observed higher PSNR values (Figure 5**b**). WU-net led to statistically significant improvements in the correlation of extracted *σ*^2^*(β)*, with GT 6X images relative to analysis of the RAW 1X images. Extracted ^*B̅*^ values were also better correlated to GT 6X images, albeit improvements were not significant.

Comparing WU-net to an identical CARE model, we observed that WU-net achieved improved performance on β metrics while maintaining recovery of RR metrics (Figure 4**c**). The overall improved β restoration suggested that WU-net was better able to capture true signals from noise in the high spatial frequencies found in NAD(P)H images. WU-net further preserved the relationship between NAD(P)H and FAD channel images, enabling high correlations for RR metrics. Due to the observed performance of WU-net for β metric recovery, we explored further optimization of WU-net which could be achieved by varying training datasets.

**Table 4:** Summary of standard metrics of image quality for RAW 1X images and denoised images generated after signal pre-processing. Values are reported for mean performance (± standard deviation) across all test set ROIs.

| Data Transform | NAD(P)H Images |  | FAD Images |  |
| --- | --- | --- | --- | --- |
| | PSNR (dB) ↑ | SSIM ↑ | PSNR (dB) ↑ | SSIM ↑ |
| **RAW 1X** | $19.2 \pm 2.8$ | $0.48 \pm 0.09$ | $23.1 \pm 5.5$ | $0.49 \pm 0.13$ |
| **CARE** | $22.8 \pm 3.0$ | **$0.63 \pm 0.08$** | **$27.2 \pm 3.7$** | **$0.62 \pm 0.07$** |
| **Wavelet** | **$23.6 \pm 2.3$** | **$0.63 \pm 0.08$** | $26.1 \pm 2.3$ | $0.57 \pm 0.09$ |

### Selection of Training Data

Initial model development focused on a limited training set of cervical tissues of known benign status (Healthy). Benign tissue samples comprised of cell layers with consistent changes in differentiation as a function of depth among image stacks. Training on such images was expected to enable the model to learn characteristics of noise without having to account for feature heterogeneity found in pre-cancerous cervical tissue samples. We further sought to assess whether training on a data set that was expanded to include image stacks from tissues with both benign and pre-cancerous lesions (Mix) impacted performance. In this new training set, 1657 and 554 RAW 1X NAD(P)H OSs (6628 and 2216 OSPs) were used for training and validation of the models, respectively. An independent test set of 1018 OSs (4072 OSPs) was used to evaluate model performance after training.

**Figure 5:**
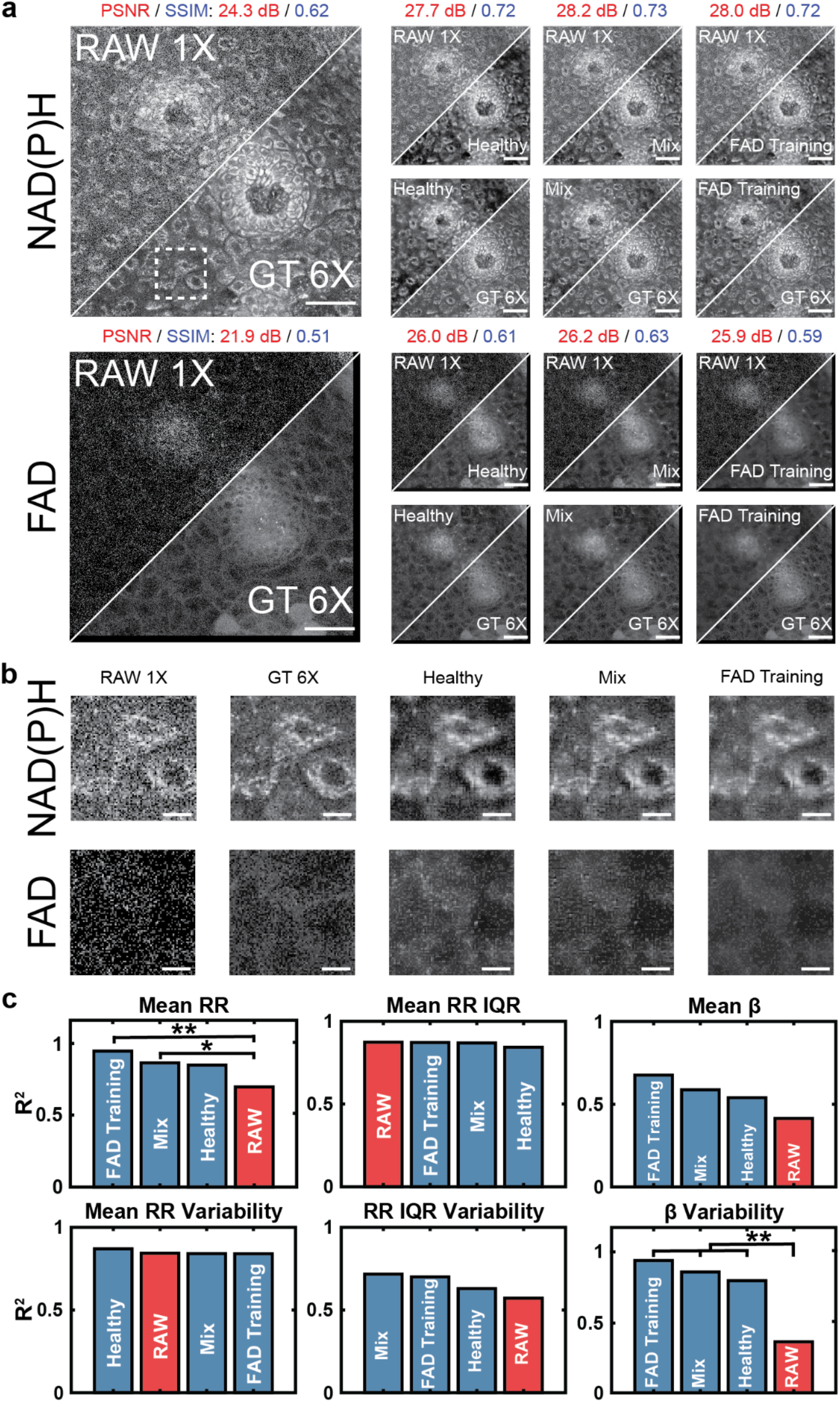
**(a)** A 290 x 290 μm^2^ field of view from a LSIL cervical tissue biopsy. NAD(P)H and FAD images for the same region are shown along with the corresponding denoised image from each data type used as training data for the WU-net model (NAD(P)H Healthy Only-, NAD(P)H Mixed Diagnosis-, and FAD Mixed Diagnosis-Wavelet transformed images). All models were equally constructed with only the data type and diagnosis type varied. Scale bar = 50 μm. **(b)** A 44.2 x 44.2 μm^2^ field of view (white square in **a**) of three cells. NAD(P)H and FAD images are shown for all data types used during training and the input and ground truth images. Scale bar = 10 μm. **(c)** Bar plots of the coefficient of determination of all downstream metrics for images denoised by models trained using varying data types and diagnosis types and RAW 1X versus the GT 6X image. Fisher r to z transformation was used to measure significance. *p<0.05 and **p<0.01

An additional consideration we explored was the impact of the source of image contrast, i.e., NAD(P)H or FAD, used for training. NAD(P)H images featured greater structural information compared to FAD images, and they were utilized in our analysis for extraction of mitochondrial clustering-focused metabolic function metrics (Figures 3-5). Thus, training was focused on NAD(P)H images, and optimized model weights from NAD(P)H image training were used to denoise FAD images for extraction of RR-based metrics. However, since similar noise characteristics were assumed to be present in both RAW 1X NAD(P)H and FAD images, we sought to confirm that training on NAD(P)H images was optimal. Thus, we used FAD images to train WU-net models using the same hyperparameters and settings as the ones used when NAD(P)H images were used. Post-training, NAD(P)H images were denoised using the weights of the FAD image trained model to extract RR and mitochondrial clustering-based metrics.

The use of training sets with mixed diagnosis images resulted in minimal differences in the denoised images when compared to training just on healthy sample images (Figure 5**a**). PSNR and SSIM values for images were observed to be nearly identical because of these insignificant differences (Table 5). Both models led to denoised images with consistent cell boundary and intracellular structures given the same RAW 1X images (Figure 5**b**) and had similar levels of restoration of downstream metrics, with the mixed diagnosis data set leading to slightly improved correlations in most cases (Figure 5**c**). The increase in correlation could be attributed to the large training set available for a mixture of diagnoses compared to only training on healthy data.

An identical model was trained using the FAD image data from the mixed diagnosis dataset. While the denoised images from the FAD-trained model looked like those from the corresponding NAD(P)H-trained model (Figure 5**a** and 5**b**), standard metrics of image quality were slightly lower. Images denoised by the FAD-trained model demonstrated higher background signal compared to images denoised by NAD(P)H-trained models (Figure 5**b**). However, despite FAD images lacking much of the structural and morphological information of their NAD(P)H counterparts, their use in training led to further improvements in β metric recovery and mean RR restoration from the RAW 1X images (Figure 5**c**). We hypothesize high frequency information in the FAD images originated primarily from noise in comparison to NAD(P)H images. As a result of the high frequency information containing primarily noise, the model improved in its’ learning of noise characteristics in the images, enabling improved denoising and recovery of metrics of metabolic activity (Figure 5**c**).

**Table 5:** Summary of standard metrics of image quality for RAW 1X images and denoised images generated after training models on various data types. Values are reported for mean performance (± standard deviation) across all test set ROIs.

|  | NAD(P)H Images |  | FAD Images |  |
| --- | --- | --- | --- | --- |
| Training Data | PSNR (dB) ↑ | SSIM ↑ | PSNR (dB) ↑ | SSIM ↑ |
| **RAW 1X** | $19.2 \pm 2.8$ | $0.48 \pm 0.09$ | $23.1 \pm 5.5$ | $0.49 \pm 0.13$ |
| **Healthy Only** | **$23.6 \pm 2.3$** | **$0.63 \pm 0.08$** | $26.1 \pm 2.3$ | **$0.57 \pm 0.09$** |
| **Mixed NAD(P)H** | $23.4 \pm 2.5$ | **$0.63 \pm 0.08$** | **$26.3 \pm 3.1$** | **$0.57 \pm 0.08$** |
| **Mixed FAD** | $23.5 \pm 2.6$ | $0.62 \pm 0.09$ | $24.8 \pm 3.7$ | $0.52 \pm 0.08$ |

### Summary of Final Model Performance

Across all models, image quality improved after denoising based on PSNR and SSIM (Table 6). Based on standard image quality metrics of all models discussed in this study, it could be assumed that models trained using NAD(P)H images and the CARE architecture with standard loss functions of MAE and MSE would perform best at the restoration of downstream metrics (Figure 6**a**). CARE models trained with MAE and MSE loss functions both demonstrated statistically significant improvement in denoised FAD and NAD(P)H image PSNR and SSIM (p<0.001). Comparatively, Wavelet-transformed-FAD images denoised using WU-net with SSIM + R2 loss had poorer standard metric performance (Table 6). Images restored with this model did not achieve statistically significant improvement of FAD image PSNR and SSIM (Figure 6**a**). As PSNR and SSIM are commonly used as indicators of model performance, it was expected that improvements in these metrics would correspond to better recovery of downstream metabolic metrics. However, the WU-net model trained on mixed diagnosis, FAD images led to denoised images whose extracted metabolic metrics were consistently correlated with the metrics extracted from GT 6X images (Figure 6**b**). The final correlations of the models shown in Figure 6 are reported in Table 7.

**Table 6:** Summary of standard metrics of image quality (PSNR and SSIM) for RAW 1X images, standard implementation of CARE, and the best performing model from this study. Values are reported for mean performance (± standard deviation).

|  | NAD(P)H Images |  | FAD Images |  |
| --- | --- | --- | --- | --- |
| Final Model | PSNR (dB) ↑ | SSIM ↑ | PSNR (dB) ↑ | SSIM ↑ |
| RAW 1X | $19.2 \pm 2.8$ | $0.48 \pm 0.09$ | $23.1 \pm 5.5$ | $0.49 \pm 0.13$ |
| Healthy NAD(P)H<br>CARE MAE | $23.6 \pm 2.8$ | $0.63 \pm 0.08$ | **$26.9 \pm 2.7$** | **$0.59 \pm 0.08$** |
| Healthy NAD(P)H<br>CARE MSE | **$23.7 \pm 2.6$** | **$0.64 \pm 0.08$** | $26.8 \pm 2.7$ | **$0.59 \pm 0.08$** |
| Mixed FAD CARE<br>SSIM + R2<br>Wavelet Transform | $23.5 \pm 2.6$ | $0.62 \pm 0.09$ | $24.8 \pm 3.7$ | $0.52 \pm 0.08$ |

## Discussion

Tissue morphological and functional metrics extracted from label-free, 2PM images could provide significant clinical utility for disease diagnosis^25^. Neural networks will likely play a critical role in enabling accurate extraction of such metrics from images that are likely to be acquired in an in vivo imaging setting. Previous studies by multiple groups have demonstrated deep learning-based denoising models can be used to improve the PSNR and SSIM of fluorescence images acquired using 2PM^33, 35, 36^. Here, we demonstrated PSNR and SSIM, while relevant in the assessment of image quality, were not representative of functional metric recovery needed for clinical utility (Figure 6).

Different algorithms have been reported for denoising of fluorescence images, however, only *Shen et al. (2022)* have reported a network used for denoising of label-free autofluorescence images^36^. In this study, a modified enhanced SRGAN model was used to denoise ex-vivo, multi-modal label-free images of human ovarian cancer tissue sections^36^. The trained GAN demonstrated an ∼4.5 dB improvement in PSNR and 79% improvement in SSIM after denoising^36^. In comparison, we demonstrated ∼4.3 dB and ∼2.7 dB improvement in PSNR and ∼30% and 6% improvement in SSIM for NAD(P)H and FAD images, respectively (Figure 6**a**). While improvement in image PSNR and SSIM were lower, RAW 1X and Denoised images in this study have higher PSNR and SSIM for all images suggesting differences in enhancement are due to limits in image improvement and not a lack of network performance (Table 6).

**Figure 6:**
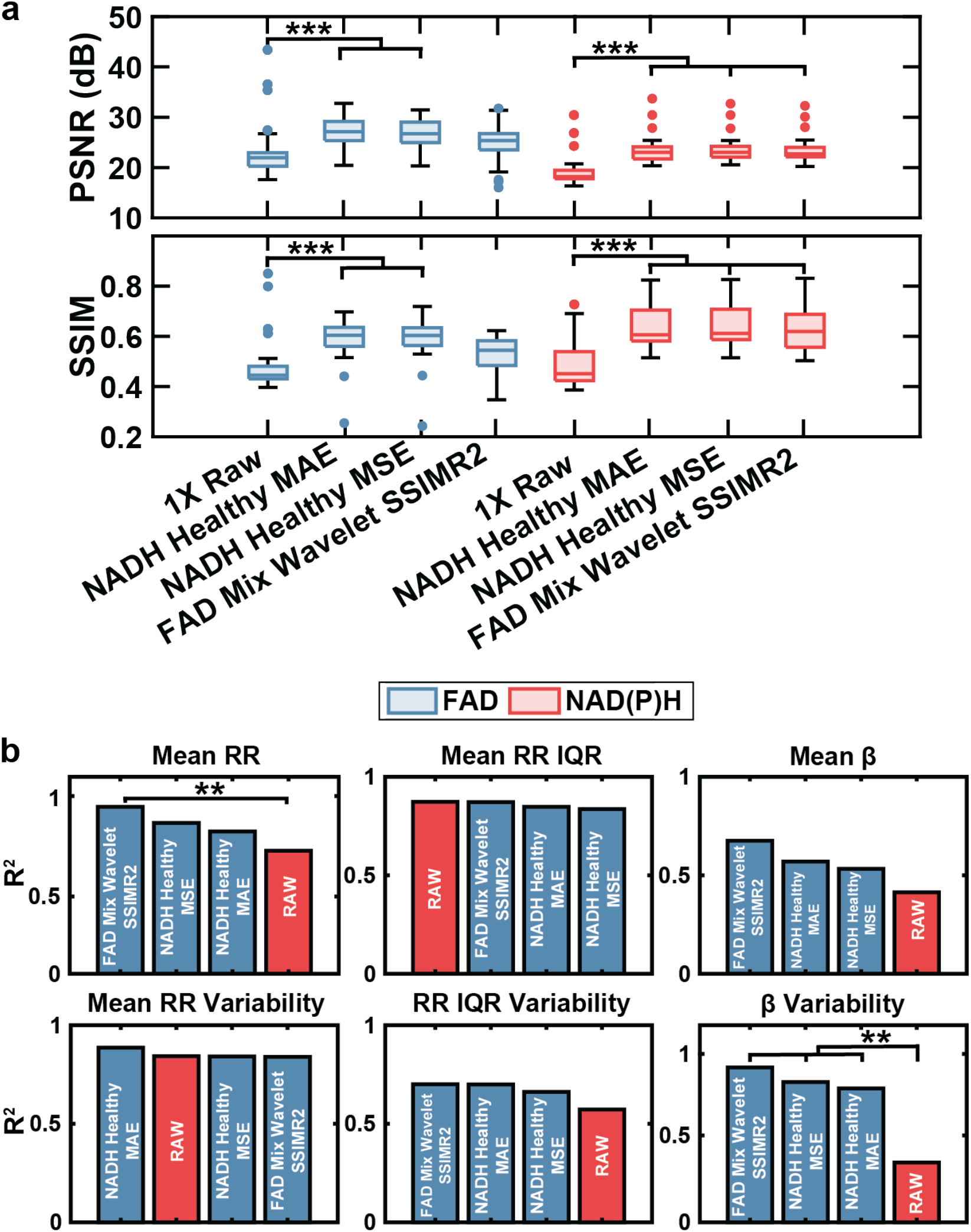
(**a**) Box and whisker plots of PSNR and SSIM of 40 test set ROIs. Denoised images demonstrated an improvement in standard metrics of image quality. (**b**) Bar plots of the coefficient of determination of all downstream metrics for images denoised by models trained using various data types, loss functions, and diagnosis types versus the GT 6X image. A one-way ANOVA with Tukey Kramer post-hoc test was used to measure significance of PSNR and SSIM. Fisher r to z transformation was used to measure significance of improvement in metabolic metric correlations *p<0.05, **p<0.01, and ***p<0.001.

**Table 7:** Correlation values of models in Figure 6. Fisher r to z transformation was used to measure significance. **p<0.01

| Final Model | Downstream Metrics |  |  |  |  |  |
| --- | --- | --- | --- | --- | --- | --- |
| | Mean RR ↑ | Mean $\beta$ ↑ | Mean RR IQR ↑ | $\sigma^2$(Mean RR) ↑ | $\sigma^2$(Mean $\beta$) ↑ | $\sigma^2$(Mean RR IQR) ↑ |
| RAW 1X | 0.71 | 0.40 | **0.87** | 0.84 | 0.33 | 0.57 |
| Healthy NAD(P)H CARE MAE | 0.82 | 0.57 | 0.85 | **0.89** | 0.78** | **0.70** |
| Healthy NAD(P)H CARE MSE | 0.87 | 0.53 | 0.84 | 0.84 | 0.81** | 0.66 |
| Mixed FAD CARE SSIM + R2 Wavelet Transform | **0.96\*\*** | **0.68** | **0.87** | 0.84 | **0.90\*\*** | **0.70** |

We further observed that GAN models did not perform well on our dataset. GANs aim to emulate characteristics of high SNR images in low SNR images through an adversarial training process^34^. To improve image quality, GANs learn the manifold of high SNR data which is assumed to be composed of images that have similar image quality metrics^44^. Thus, it is important for image quality to be consistent across all high SNR images.

High-SNR images from a single depth in a thin OS, like those used to train the enhanced SRGAN model in *Shen et al.* (2022), have similar image quality for all ground truth images leading to improved GAN performance^36^. In our study, bulk tissues were imaged at multiple depths leading to inconsistent image quality in our ground truth images as SNR is known to change as a function of depth. As such, we hypothesized the GAN model implemented in this study failed to learn the manifold of high SNR images and improve our images whereas the enhanced SRGAN model implemented by *Shen et al.* (2022) succeeded.

While multiple studies demonstrate models capable of improving PSNR and SSIM, assessment of morphofunctional metrics of metabolic activity after denoising has not been examined previously^33–37^. Here, we calculate restored image PSNR and SSIM along with metabolic metric recovery and observe that higher PSNR and SSIM values did not ensure the greatest restoration of RR and β metrics (Figure 6). While PSNR and SSIM values between models are observed to be within <5% of each other (Table 3-5), many studies indicate maximum improvement of PSNR and SSIM values as indicators of model performance^33–37^. In this study, we observe that models with optimal PSNR and SSIM values did not yield the greatest recovery of metabolic metrics. Altogether, PSNR and SSIM are not well suited for assessment of model performance on label-free 2PM images, necessitating further validation using metrics of metabolic activity.

Application of denoising algorithms on label-free 2PM datasets to date have been limited by the lack of available large clinical datasets^36, 45^. Deep learning models have shown promise with small datasets (*Shen et al.* (2022) used only 24 paired images) in image restoration; however, larger datasets are needed for consistent reconstruction of high-SNR images^36, 44^. Here we presented a denoising network trained on a larger training set of 1657 OSs (6628 OSPs) and evaluated on an independent test set of 1018 OSs (4072 OSPs).

Using CARE, we observed improvements in image quality based on standard metrics (Table 3). However, the pre-packaged, standard models showed poor recovery of β metrics. Custom-loss functions improved metabolic metric recovery by penalizing models for both failing to generate similar images and reducing pixel correlation (see Supplementary Table S2 online). More interestingly, we observed the use of DWT to separate the frequency information in an image before training independent models (WU-net) produces images that had high metabolic metric correlations with GT 6X metrics (Figure 4**c**). By training on independent frequency-band images, the models were forced to learn the noise characteristics of different frequency bands without convolving the bands^42^.

A key advantage of WU-net, in comparison to identically trained (non-wavelet) U-nets, was the denoising of higher frequencies where noise was expected to be dominant. Denoising of high frequency noise led to enhanced recovery of β metrics as WU-net was more consistent in reducing noise in these frequencies (See Supplementary Discussion S3 and Supplementary Fig. S3 online). WU-net led to a statistically significant decrease in high frequencies compared to a comparable CARE model (See Supplementary Fig. S3 online). Further, the incorporation of SSIM + R2 as a loss function promoted the models to restore similar frequencies from the GT 6X image in the denoised image while minimizing the loss of correlation between pixels.

Further, we observed that models trained on FAD images outperform their NAD(P)H counterparts (Figure 5**c**). To explain this phenomenon, we examined the correlation of optical RR metrics between RAW 1X images and GT 6X images. RR metrics from RAW 1X images correlated well with RR metrics from GT 6X images, suggesting that the noise characteristics in FAD and NAD(P)H images are similar. However, as the FAD images contain less signal compared to their NAD(P)H paired images, high-frequency contributions are mostly noise in the RAW 1X FAD images. Thus, training on FAD images likely improved the model’s learning of noise characteristics. This led to improvement in downstream metric recovery and translation of model weights to NAD(P)H images.

WU-net with a custom loss (SSIM + R2) function and training on FAD data demonstrates improved restoration of most metrics of metabolic activity from label-free, 2PM images (Table 7); however, further improvements in the restoration of ^*B̅*^ are desired. One potential method of improving ^*B̅*^ restoration would be to design a loss function that minimizes the differences in the power spectral density maps of paired images that are used for β calculation. A challenge of such a method would be the computational time required for generating these maps^22, 24, 25^. Future studies may examine simpler predictors of mitochondrial clustering using a modified GAN network, where the discriminator network will estimate β from the input images and optimize the generator to achieve accurate β metric recovery.

In this study, we specifically focused on restoration of morphological and functional metrics from label-free, 2PM images of human cervical tissue, relying on a single denoising algorithm. Future studies will examine the application of the trained denoising model and model architecture on datasets acquired from different microscope systems, objective lenses, and tissue types. Validation of the model on these datasets would support the broad use of WU-net for denoising label-free 2PM images. Further, successful implementation of pre-trained models on other datasets would reduce the need for large clinical datasets^36, 45^. As the model advances, improvements in ground truth data collection are needed. Ground truth data used in this study contain noise and therefore are not truly representative of mitochondrial signal. Alternative techniques for image acquisition such as slower line scan speed could be utilized to improve ground truth image quality.

In summary, we demonstrated that maximizing standard metrics of image quality (PSNR and SSIM) did not necessarily lead to improved recovery of functional tissue metrics, especially ones associated with mitochondrial organization (Table 7). Using WU-net with a custom loss function, we demonstrated improved recovery of functional metrics of metabolic activity, even though PSNR and SSIM metrics were not optimal. Results from this study support the application of deep learning algorithms for the restoration of RR and β metrics from low-SNR 2PM images. As more data becomes available both from varying microscope systems, objective lenses, and tissue types, a more robust algorithm could be generated for rapid image collection and classification, eventually improving patient health during *in vivo* image collection.

## Materials and Methods

### Sample Acquisition

All activities pertaining to cervical tissue biopsy handling were done in accordance with approved Tufts Health Sciences IRB protocol #10283. Patients over the age of 18 with a recent LSIL or HSIL pap smear diagnosis undergoing colposcopy or loop electrosurgical excision procedure (LEEP) were recruited to the study. Informed consent was acquired from all study subjects before participation. During the routine procedure, a second biopsy from a colposcopically abnormal region of the cervix was taken and placed in a custom-built tissue carrier containing keratinocyte serum-free media (KSFM; Lonza). Biopsies were transported via personal vehicle to the Tufts Advanced Microscopy Imaging Center (TAMIC) for imaging. All imaging was conducted within 4 hours post-biopsy. Immediately after imaging, biopsies were fixed in 10% neutral buffered formalin. Biopsies were returned within 5 business days to the Tufts Medical Center Department of Pathology for standard histopathological diagnosis.

Patients over the age of 18 undergoing hysterectomies for benign gynecological disease were also recruited to the study as healthy controls. The only difference between healthy and precancerous biopsy acquisition was in the actual biopsy excision. Healthy biopsies were sampled from the resected cervix by a pathologist after macroscopic inspection to rule out abnormalities.

### Deep Learning Dataset Details

A total of 151 ROIs (image stacks) were collected from 54 patients. The training and validation set was comprised of 100 ROIs featuring 5-50 OSs per ROI. 75% of the ROIs were randomly selected for training and the remaining 25% were set aside as a validation set (1657 training OSs and 554 validation OSs). The test set featured 51 ROIs (with 10-50 OSs per ROI) and was excluded from all training (1018 OSs). For *k-*fold validation, training and validation sets were shuffled for up to five seeds to ensure robustness of denoising on a constant test set (see Supplementary Fig. S6 online). The dataset features images from tissues with three diagnoses: Benign, LSIL, and HSIL. The test set was composed of 25 Benign ROIs (49.02%), 14 LSIL ROIs (27.45%), and 12 HSIL ROIs (23.53%). The training and validation sets were composed of 55 Benign ROIs (54.45%), 25 LSIL ROIs (24.75%), and 21 HSIL ROIs (20.79%). Based on training/validation splitting seed, these values could range from 52-57.3% Benign, 25.3-26.7% LSIL, and 18.7-22.7% HSIL in the training set and 48-64% Benign, 20-24% LSIL, and 16-28% HSIL in the validation set. An alternative training scheme was initially attempted. In this scheme, only benign ROIs were used in training with 112 ROIs of mixed diagnosis being used in the test set and 39 benign ROIs being used for training. The training set was later modified as it became evident that training on a mixture of diagnoses resulted in superior restoration of downstream metrics (Figure 5).

### Optical Instrumentation and Image Acquisition

Images were collected using a commercially available Leica SP8 inverted microscope system equipped with an Insight fs laser. Tissue biopsies were placed epithelial side down onto a glass bottom dish and light was delivered using an epi-illumination scheme. Tissue biopsies were excited with 755 nm and 860 nm light. Two hybrid photodetectors (HyDs) were set up to collect 460 ± 25 and 525 ± 25 nm light. Two photomultiplier tubes (PMTs) were set up to collect 430 ± 12 and 624 ± 20 nm light.

Light was delivered and collected using a 40X/1.1 NA water-immersion objective lens (290 x 290-μm field-of-view). Images were collected through the full thickness of the epithelium using a depth-sampling rate of 4-μm. Six individual frames were collected at each depth. On average, 3 – 5 ROIs were sampled from each biopsy.

### Morphological and Functional Metrics

Images were calibrated and processed as described in detail previously to extract images that represented NAD(P)H and FAD TPEF intensity fluctuations^23–25, 46^ . At each optical depth, NAD(P)H and FAD images were used to define a corresponding redox ratio for each pixel of the field, as:

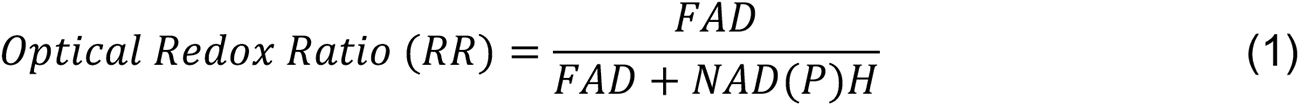

From the RR distributions for each OS, we calculated the mean RR and the interquartile range (IQR) as metrics of the overall oxidation-reduction tissue state and the corresponding heterogeneity, respectively. The mean and sample variance (variability) of the mean OS RR and the OS RR IQR for all images in an epithelial stack were calculated to assess the depth-dependence of these metrics.

NAD(P)H images were analyzed as described previously ^17, 18, 21, 22^ using a Fourier based approach to extract a value for the parameter β, as a metric of the level of mitochondrial fragmentation and networking, which also depends highly on the metabolic activity of the tissue. Briefly, an inverse power law was fit to the power spectral density (PSD) of the 2D Fourier transform of the cytoplasmic NAD(P)H intensity fluctuation images, as:

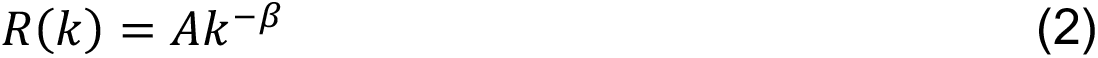

where *R* is the fit to the PSD, *k* is the magnitude of the spatial frequency, *β* is the power law exponent, and *A* is a constant. The mean and sample variance of *β* were assessed as a function of depth for each image stack.

### Deep Learning Model Description

The basic structure of the CARE network has been described extensively (See Supplementary Fig. S4**a** online)^33^. The network was implemented through Keras and TensorFlow^47, 48^. A copy of the CSBDeep repository (https://github.com/CSBDeep/CSBDeep) was locally imported into an anaconda environment^49^. The network was configured to take a 256 x 256 x 1 input image and generate a 256 x 256 x 1 denoised image. A 40-gigabyte Nvidia Tesla A100 GPU card was used for all training and evaluation. Typically, a 1 x 512 x 512 x z-depths image stack was split into 4 x 256 x 256 x z-depths image patches before training. A starting learning rate of 1 x 10^-5^ was used with an Adam optimizer^50^. Training was allowed to continue for 300 epochs with a scheduler reducing the learning rate when network performance stagnated for more than 20 epochs. Loss functions were varied to find the optimal function to improve downstream analysis performance. Loss functions used include SSIM Loss, R2 Loss, Focal Frequency Loss (FFL), MAE (L_1_) Loss, MSE (L_2_) Loss, and combined losses such as a combined SSIM + L_2_, SSIM + FFL, SSIM + R2 Loss^51^. Six down-sampling and up-sampling layers were generated with the first layer expanding the single-channel images to thirty-two channels. Residual connections were used to preserve encoded information from each down sampled layer and pass it forward to the decoder layers (see Supplementary Figure S4 online).

For the WU-net architecture, four CARE networks, one per subband, were built as described above. A DWT was used to decompose a 1 x 256 x 256 OSP into 1 x 128 x 128 x 4 frequency band images. The four frequency bands would then be individually input to each CARE network for denoising. After denoising, an IDWT was used to reconstruct the 1 x 256 x 256 OSP (for greater detail see Supplementary Fig. S5 online).

Training time typically varied from one to two hours, with an evaluation time of approximately twenty-four seconds per image stack. For all trained CARE networks, 3D SSIM, PSNR, Mean β, β Variability, Mean RR, RR Variability, RR IQR, RR IQR Variability were analyzed. All final metrics were assessed using a single frame input, denoised, and ground truth (6 frame averages) images with built-in and custom MATLAB (MathWorks; Natick, MA) functions.

### Statistics

For Figures 3-5**c** and Table 7, Fisher r-to-z transformation was used to convert Pearson’s correlation coefficients (r) to z_r_ values^52^. This transform was calculated using Equation 3:

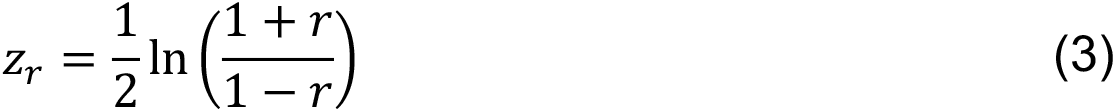

The z_r_ value, unlike r, belongs to a normal distribution, allowing for the calculation of a Z-statistic to determine confidence intervals. To calculate the test Z-statistic for comparison of zr values to determine significance, Equation 4 was used:

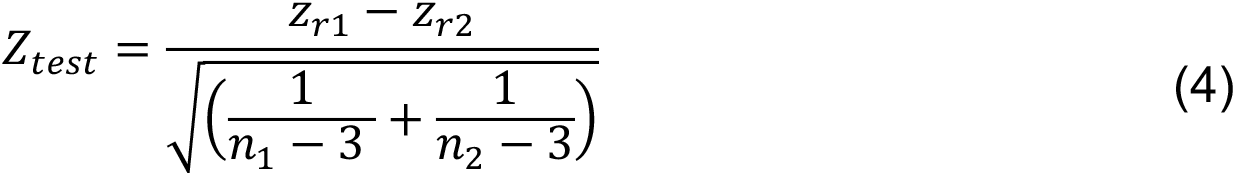

where n_1_ and n_2_ are the sample size of r_1_ and r_2_, respectively^53^. The Z_test_ value was then compared to the critical Z-values to determine significance and p-values using a two-tailed distribution.

## Data Availability

The raw datasets used for model generation in the current study along with the trained model weights are available from the corresponding author on reasonable request.

Codes for network training and prediction are publicly available at https://gitlab.tufts.edu/georgakoudi-lab/Denoising2PImages

## Acknowledgments

We would like to thank the National Institute of Biomedical Imaging and Bioengineering (R01 EB030061), the National Institute of Health, Office of the Director (S10 OD021624), and the National Cancer Institute for funding this work (R03 CA235053).

The authors acknowledge the Tufts University High Performance Compute Cluster (https://it.tufts.edu/high-performance-computing) which was utilized for the research reported in this paper. The support of the Data Intensive Center (DISC) is acknowledged. We would also like to thank Jasmine Kwan for her help in preparing figures for this paper.

## Author Contribution

I.G. conceived the initial goals of the study. Under the guidance of I.G., C.P. collected all tissue biopsies used in this study and subsequently collected the image datasets. H.T. screened and recruited patients for this study. E.G. and N.J. rendered biopsy diagnoses after imaging was completed. N.V. wrote the initial implementations for denoising and with F.S. further advanced the algorithms and developed a repository used to denoise the collected images. C.P. aided in modifying existing downstream analysis scripts for application on denoised images and completed statistical analysis on downstream metrics. G.G. provided guidance in code implementation and data augmentation. E.M., A.P., G.G., I.G., led discussions on interpretation of results and methods for further optimization of generated models. I.G. supervised the study and with N.V. and C.P. prepared the manuscript text. All authors have reviewed and approved the manuscript.

