## Supplementary for "Restoration of metabolic functional metrics from label-free, two-photon cervical tissue images using multiscale deep-learning-based denoising algorithms"

Supplementary Discussion S1: While loss functions such as mean absolute error (MAE or $L_1$) and mean squared error (MSE or $L_2$) are often used in other denoising studies, they face limitations on certain images due to the assumption of pixel-wise independence which may not be true in all cases[1–3]. To improve image similarity, a SSIM loss function was implemented to minimize image dissimilarity between the denoised and GT 6X images. While promising, blurring and a loss of high frequency variance was observed in denoised images because of the Gaussian filter (Supplementary Fig. S1). Various filter sizes and sigma values were evaluated prior to selection of a 3 x 3 filter (Supplementary Table S1). Inclusion of pixel-wise loss functions in addition to SSIM loss was hypothesized to enable high visual similarity in denoised images while also preventing the loss of high frequency variance in images. Thus, in addition to MAE ($L_1$) and MSE ($L_2$), we tested three additional loss functions that penalized pixel-wise differences in combination with the SSIM loss function: SSIM + $L_2$, SSIM + the coefficient of determination (R2), SSIM + Frequency Focal Loss (FFL) (Supplementary Fig. S2)[4]. FFL utilizes Fast-Fourier Transforms (FFT) to calculate the frequency map of each denoised and GT 6X image and penalizes the difference in phase and magnitude between the two images[4]. The impact of these five loss functions on denoising was assessed using the CARE model.

MAE, MSE, and SSIM + FFL all led to the generation of visually similar images, yielding similar metrics of image quality (Supplementary Fig. S2**a**). SSIM + R2 and SSIM + L2 loss functions led to weaker image quality metrics, albeit differences were insignificant across the entire test set (Supplementary Table S2). Differences in image quality were associated with recovered signal from the nucleus and interstitial space with smaller fluctuations in the cytoplasm of cells which varied between loss functions (Supplementary Fig. S2**b**). The small fluctuations in cytoplasmic signal were of particular interest as the calculation of metabolic metrics was associated with NAD(P)H and FAD intensity measurements from the cytoplasm.

While standard metrics of image quality suggested all loss functions yielded similar images, differences in cytoplasmic signal accounted for varying performance on downstream metrics. Images restored by MSE, MAE, and SSIM + FFL all led to statistically significant improvements in β variability correlation between the denoised and GT 6X values (Supplementary Fig. S2**c**). This was consistent with the improved PSNR of restored NAD(P)H images (Supplementary Table S2). MAE and MSE loss both generated FAD and NAD(P)H images with high PSNR values, leading to 1-5% improvements in recovery of some of the RR metrics. SSIM + R2 was the only loss function to demonstrate statistically significant recovery of RR IQR variability. In comparison to other loss functions, SSIM + R2 loss led to improved FAD image PSNR; however, NAD(P)H image PSNR was poor (Supplementary Table S2).

In principle, the loss functions with the higher PSNR and SSIM values (Supplementary Table S2) led to the recovery of metabolic function metrics that were highly correlated with GT 6X images. However, no loss function led to statistically significant improvement in mean β ($\bar{\beta}$).

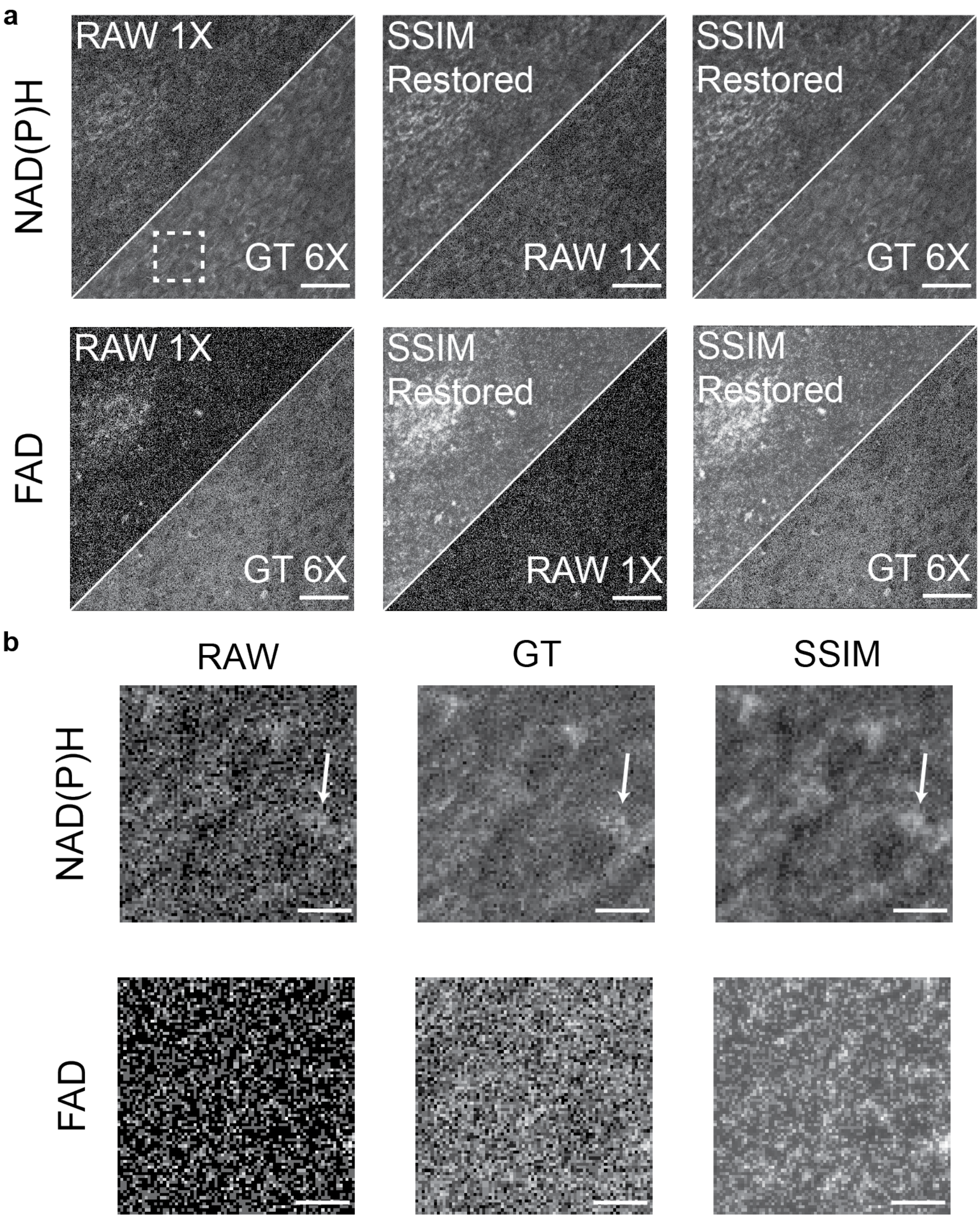


Supplementary Figure S1: **(a)** A 290 x 290 μm$^2$ field of view from a benign cervical tissue biopsy. NAD(P)H and FAD images for the same region are shown along with the corresponding denoised image after using an RCAN model with SSIM loss. Scale bar = 50 μm. **(b)** A 44.2 x 44.2 μm$^2$ field of view (white square in a) of two cells. NAD(P)H and FAD images are shown to demonstrate the blurring effect observed when using SSIM loss for denoising. The white arrows demonstrate how high intensity regions are blurred in the denoised image leading to changes in the spatial frequency distribution of the image. Scale bar = 10 μm.

Supplementary Table S1: Correlation values of WU-net models trained using SSIM + R2 loss with varying gaussian filter sizes and sigma values. Fisher r to z transformation was used to measure significance. **p<0.01

| | Downstream Metrics | | | | | |
|---|---|---|---|---|---|---|
| **FAD WU-net SSIMR2 Loss** | **Mean RR ↑** | **Mean β ↑** | **Mean RR IQR ↑** | **$\sigma^2$(Mean RR) ↑** | **$\sigma^2$(Mean β) ↑** | **$\sigma^2$(Mean RR IQR) ↑** |
| **RAW 1X** | 0.71 | 0.43 | **0.87** | 0.84 | 0.22 | 0.57 |
| **Filter Size = 3, σ = 0.5** | **0.96**** | 0.68 | **0.87** | **0.84** | **0.90**** | 0.70 |
| **Filter Size = 7, σ = 1.0** | **0.96**** | **0.69** | 0.84 | 0.77 | 0.85** | 0.66 |
| **Filter Size = 11, σ = 1.5** | **0.96**** | 0.65 | 0.86 | 0.77 | 0.83** | **0.72** |

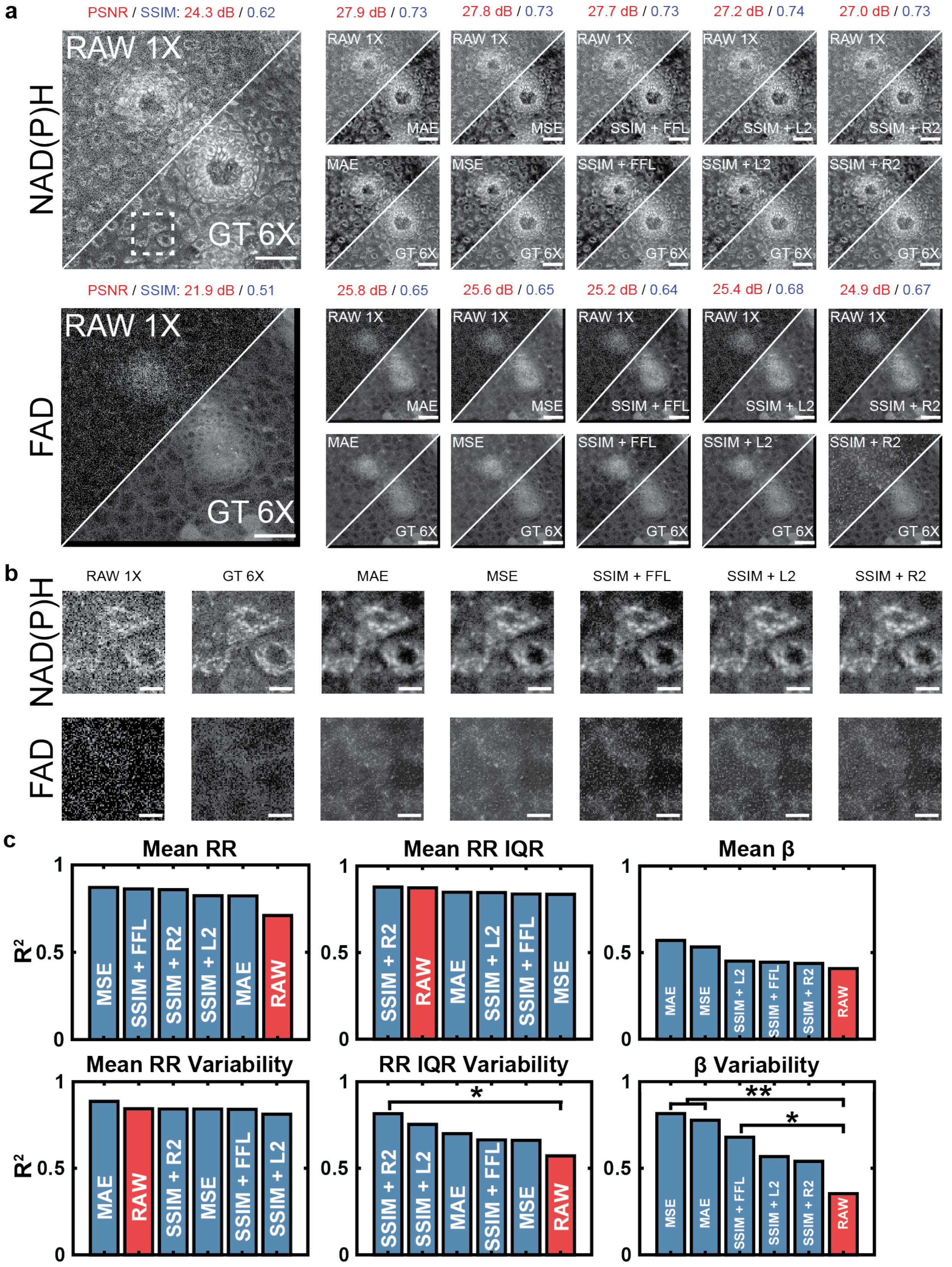


Supplementary Figure S2: **(a)** A 290 x 290 μm$^2$ field of view from a LSIL cervical tissue biopsy. NAD(P)H and FAD images for the same region are shown along with the corresponding denoised image from each objective function utilized with the CARE model (MAE, MSE, SSIM + FFL, SSIM + L2, and SSIM + R2). Scale bar = 50 μm. **(b)** A 44.2 x 44.2 μm$^2$ field of view (white square in **a**) of three cells. NAD(P)H and FAD images are shown for each CARE model based on the corresponding loss function used during training and the input and ground truth images. Scale bar = 10 μm. **(c)** Bar plots of the coefficient of determination of all downstream metrics for images denoised by CARE models trained based on the corresponding loss function during training and RAW 1X vs. the GT 6X image. Fisher r to z transformation was used to measure significance. *$p<0.05$ and **$p<0.01$.

Supplementary Table S2: Summary of standard metrics of image quality for RAW 1X images and denoised images generated from various loss functions with CARE. Values are reported for mean performance (± standard deviation) across all test set ROIs.

| | NAD(P)H Images | | FAD Images | |
|---|---|---|---|---|
| **Loss Function** | **PSNR (dB) ↑** | **SSIM ↑** | **PSNR (dB) ↑** | **SSIM ↑** |
| **RAW 1X** | 19.2 ± 2.8 | 0.48 ± 0.09 | 23.1 ± 5.5 | 0.49 ± 0.13 |
| **MAE** | 23.6 ± 2.8 | 0.63 ± 0.08 | 26.9 ± 2.7 | 0.59 ± 0.08 |
| **MSE** | **23.7 ± 2.6** | **0.64 ± 0.08** | 26.8 ± 2.7 | 0.59 ± 0.08 |
| **SSIM + FFL** | 23.6 ± 2.6 | **0.64 ± 0.08** | 26.7 ± 2.6 | 0.59 ± 0.09 |
| **SSIM + $L_2$** | 22.7 ± 2.9 | 0.63 ± 0.08 | 26.8 ± 3.1 | 0.60 ± 0.07 |
| **SSIM + R2** | 22.8 ± 3.0 | 0.63 ± 0.08 | **27.2 ± 3.7** | **0.62 ± 0.07** |

Supplementary Discussion S2: 2D-DWT is a mathematical technique utilized to separate spatial frequency information from images. Mother wavelets are applied to extract an “approximate” (LL) and detailed images (HL, LH, HH) from an image. The detailed images are comprised of horizontal (HL), vertical (LH), and diagonal (HH) features from the original image. Each of these four images (LL, HL, LH, and HH) have dimensions that are 50% of the original image, i.e., a 256 x 256 RAW 1X image will output 4, 128 x 128 images. To implement WU-net without convolving the frequency bands, four successive models were independently trained to denoise each frequency band before restoring the denoised image (Supplementary Figure S5). In this implementation, the denoising of images was found to be faster as inputted frequency images (LL, HL, LH, and HH) are a fraction of the size of the original images. Once denoised, images undergo inverse discrete wavelet transformation to restore the image to the original dimensions. For loss calculation, each frequency band output is compared to the corresponding frequency band from the ground truth image. This allows each frequency band model to optimize weights independent of the other frequency bands. The cumulative model is therefore believed to optimally denoise high and low frequency noise in the images.

Supplementary Table S3: Correlation values of WU-net models trained using NAD(P)H data and varying loss functions. Fisher r to z transformation was used to measure significance. *p<0.05, **p<0.01, ***p<0.001

| | Downstream Metrics | | | | | |
|---|---|---|---|---|---|---|
| **NAD(P)H Data WU-net** | **Mean RR ↑** | **Mean β ↑** | **Mean RR IQR ↑** | **$\sigma^2$(Mean RR) ↑** | **$\sigma^2$(Mean β) ↑** | **$\sigma^2$(Mean RR IQR) ↑** |
| **RAW 1X** | 0.71 | 0.43 | 0.87 | 0.84 | 0.22 | 0.57 |
| **SSIM + R2 Loss** | 0.88* | 0.59 | 0.87 | 0.84 | **0.82*****| 0.71 |
| **SSIM + FFL** | **0.92**** | **0.60** | **0.90** | **0.89** | **0.82***** | **0.72** |
| **MAE Loss** | 0.88* | 0.59 | 0.86 | 0.87 | 0.75** | 0.66 |
| **MSE Loss** | 0.87* | 0.47 | 0.88 | 0.84 | 0.73** | 0.64 |

Supplementary Discussion S3: WU-net based denoising demonstrates improved recovery of β metrics compared to identical CARE models. To understand this phenomenon, a closer examination of power spectral density (PSD) curves and curve-fits were examined. PSD vs. frequency plots (Figure S4**a**) were generated for all z-depths across all 51 test set ROIs. Curve-fitting was completed to extract the mitochondrial clustering power law fit. PSD values at the start and end frequencies of the power law fit were stored for downstream analysis. An array of PSD values was generated for both the WU-net (Wavelet) and CARE (Non-Wavelet) denoised images. PSD values were compared between both models to determine which frequencies were being denoised by WU-net in comparison to CARE. WU-net did not demonstrate any significant difference in the PSD values compared to CARE at the lower range of high spatial frequencies. However, WU-net significantly reduced the PSD value at the highest discrete spatial frequencies of the mitochondrial clustering power law fit in comparison to CARE models ($p<0.05$). The denoising of the highest spatial frequencies by WU-net demonstrated consistent performance, resulting in more reliable β metric recovery (Figure S4**b**). In comparison, CARE models demonstrated greater variability which in turn compromised β metric recovery. Together, these results indicate that wavelet-based denoising provides greater denoising of high frequency noise, which improves the consistency of β metric restoration.

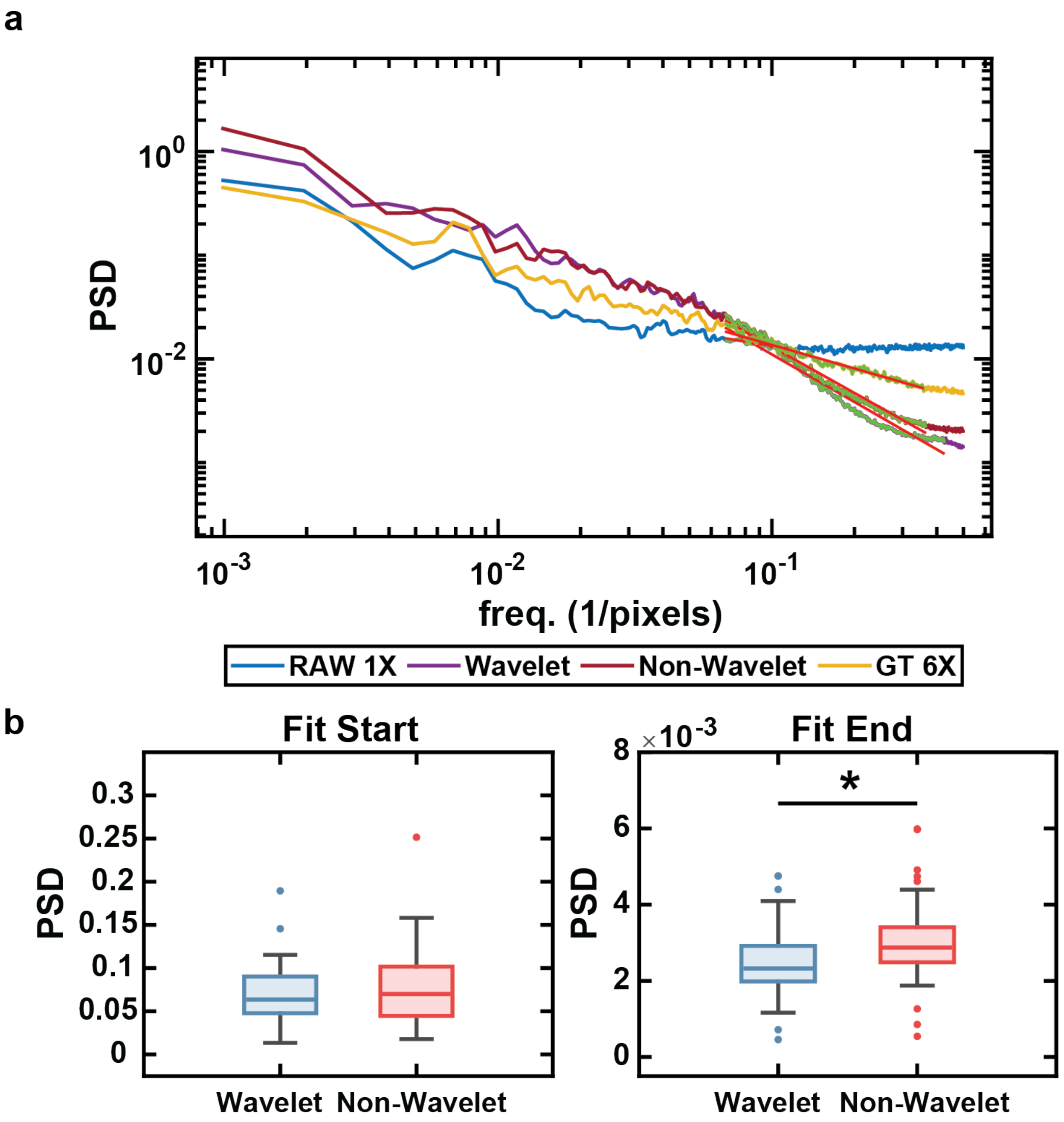


Supplementary Figure S3: **(a)** A sample power spectral density curve for a cervical tissue image. Curve fitting is used to calculate the slope of the power spectral density for sub-cellular features. Images denoised by WU-net (Wavelet) demonstrated lower PSD values for high frequencies while images denoised by an identical CARE (Non-Wavelet) demonstrated higher PSD values at high frequencies. **(b)** Characterizing the difference in PSD values at the start and end of curve-fitting, both Wavelet and Non-Wavelet demonstrated insignificant differences in PSD values where curving fitting starts but statistically significant differences in PSD where fitting ends (p-value = 0.0105). Statistical differences were calculated using a two-tailed t-test. *p<0.05.

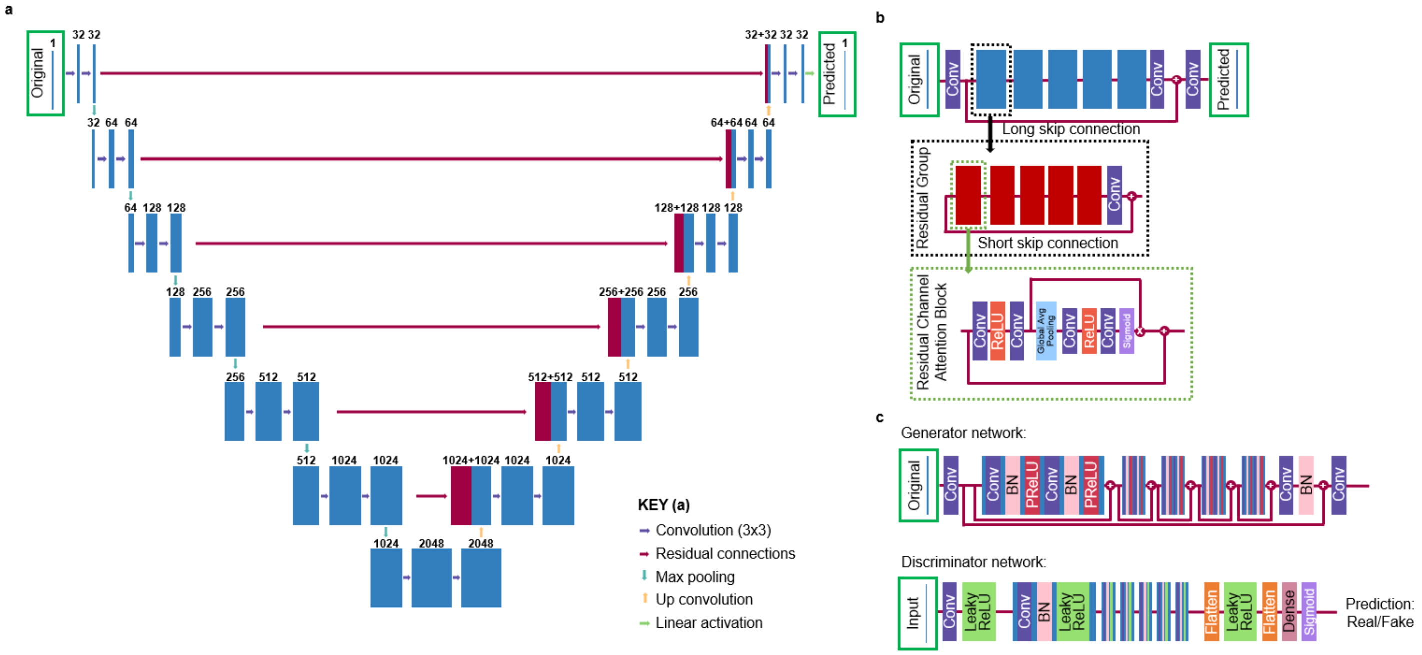


Supplementary Figure S4: **(a)** Reproduction of CARE architecture previously utilized in other studies with demonstrated residual connections and corresponding model depth from this study[1,5]. **(b)** Reproduction of the RCAN architecture initially discussed by *Zhang et al.* and later modified and implemented by *Chen et al.*[2,6]*.* **(c)** Reproduction of SRGAN model utilized by *Ledig et al.*[7].

.

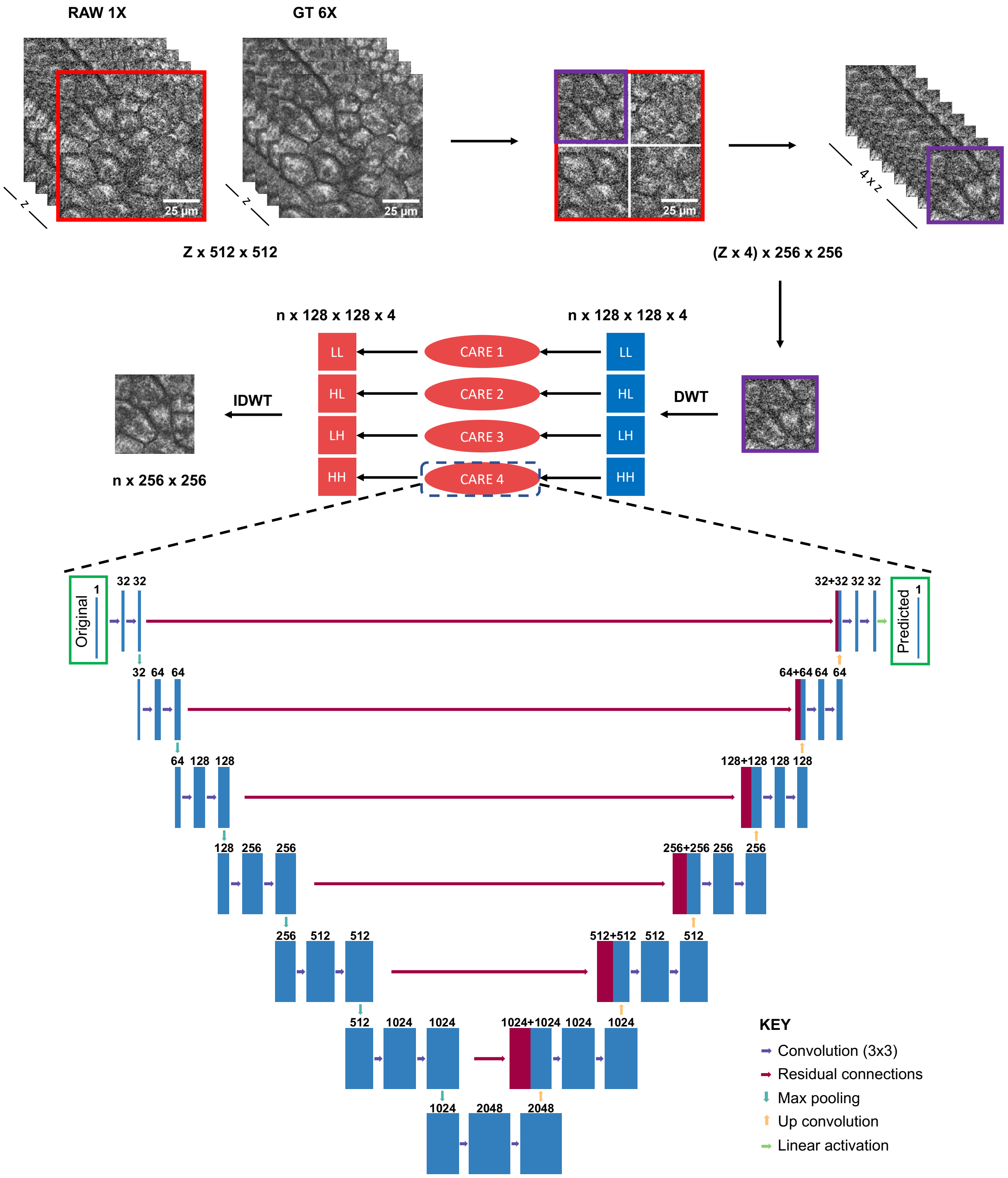


Supplementary Figure S5: Summary of the data preprocessing and model implementation pipeline for WU-net. Z x 512 X 512 image stacks from single frame and ground truth datasets are split into (4 x Z) x 256 x 256 image sets. Each 256 x 256 image is decomposed into frequency bands using a DWT generating n x 128 x 128 x 4 images where n is equal to 4 x Z. Each frequency band is passed through an identical CARE network for denoising. Denoised images are restored back into a n x 256 x 256 using an inverse DWT (IDWT).

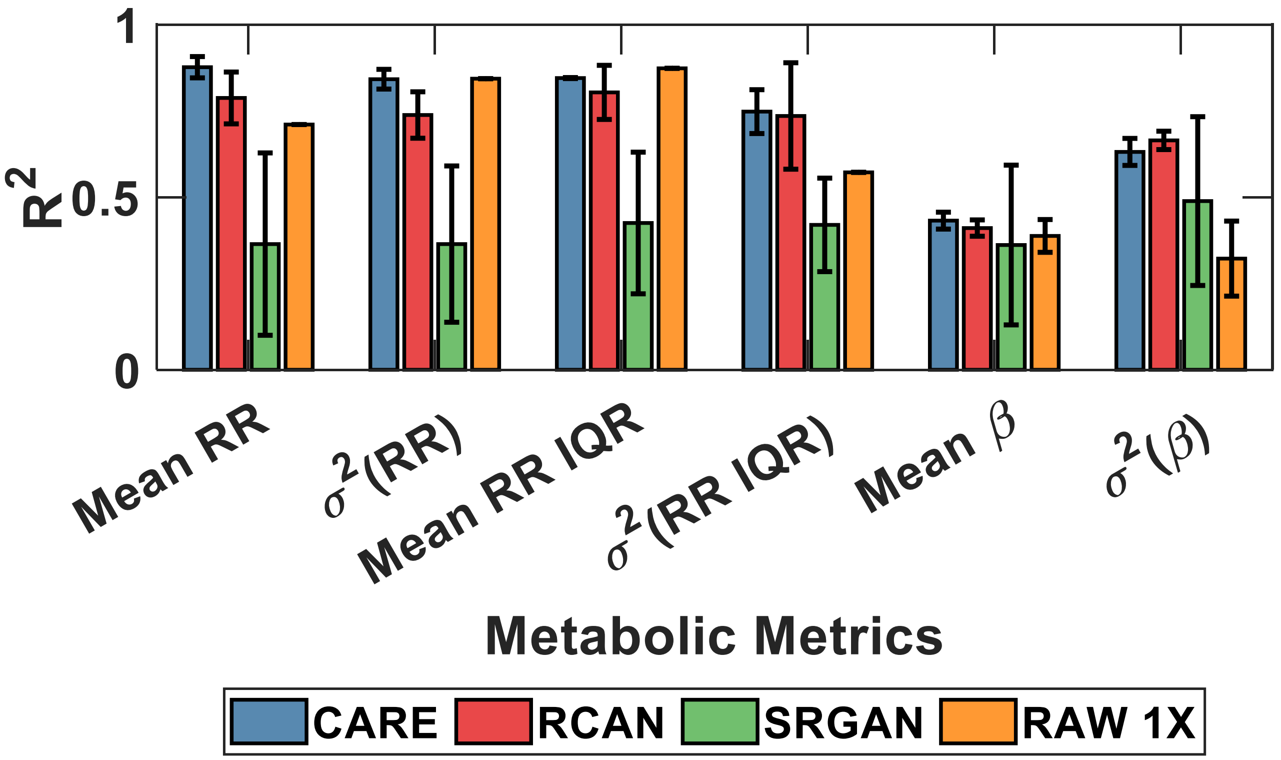


Supplementary Figure S6: Grouped bar plots of the mean coefficient of determination for the metabolic metrics extracted from images denoised using the CARE, RCAN, and SRGAN architectures and the RAW 1X images compared to metrics extracted from the GT 6X images. All models were trained using NAD(P)H data from healthy samples and SSIM + L2 loss. Mean coefficients of determination are calculated from *k*-fold validation on an independent test set using a *k* = 5. Error bars represent one standard deviation from the mean performance for each model.

**Supplementary Methods:**

*Deep Learning Performance Benchmark*

To benchmark the model performance, alternative models were trained using the same hyperparameters and datasets. RCAN was first described by Zhang et al. (2018) and later expanded for denoising of 3D datasets by Chen et al. (2021)[2,6]. The basic architecture of RCAN features multiple residual groups (RG) in which multiple residual channel attention blocks (RCAB) exist (Supplementary Fig. S4**b**). RCAN utilizes long- and short-skip connections to bypass low spatial frequencies and emphasize high spatial frequency differences. A copy of the 3D-RCAN repository (https://github.com/AiviaCommunity/3D-RCAN) was locally copied and altered to enable interfacing with other model architectures and loading wrapper functions. The trained architecture utilized five residual groups composed of five residual channel attention blocks. For feature extraction and residual layers within the RG and RCAB, the desired number of filters (channels) was set to thirty-two. Within the RCAB, a convolutional layer downscaled the thirty-two filters by a factor of four, generating eight filters. In line with modifications by Chen et al. (2021), the upscaling module from the original RCAN model by Zhang et al. (2018) was omitted as input and output images were the same size[2,6]. For consistency, all images were patched identically for all trained models as described above. Training time typically varied from four to five hours, with an identical evaluation time as the CARE model.

SRGAN is an adversarial network composed of a residual neural network (ResNet) and a simple convolutional neural network (CNN) (Supplementary Fig. S4**c**)[7]. The ResNet functions as a generator network, using a series of residual skip connections, to convert low SNR images into a predicted high SNR image[7,8]. The predicted image along with paired ground truth images were then used to train a CNN to classify generated images from ground truth images. The objective of the generator model was to predict images that were realistic and mislead the discriminator network, while the discriminator network's objective was to correctly separate the predicted and ground truth images[7].

By playing this "game" iteratively, the ResNet will eventually learn how to mislead the CNN and can be trusted to generate images that match high SNR images. To train such a model using a single GPU core, the ResNet depth was reduced to five residual-in-residual dense blocks from the standard sixteen blocks. The number of residual-in-residual dense blocks remains consistent with the number of RCABs utilized in the RCAN model[2].

Zhang et al. (2018) describes a novel perceptual loss using high-level feature maps of a well characterized Visual Geometry Group (VGG) network which was trained on three-dimensional, colored images[7,9]. However, as our dataset consists of two-dimensional, grayscale images, the VGG network was replaced with a CARE network with pre-trained weights. [9] In SRGAN, high-level features were extracted from a convolution layer within the VGG network[7]. By substituting the trained CARE model for the VGG network, the model still extracts the features needed to calculate perceptual loss with an architecture already configured for the dataset. SRGAN demonstrated improved performance when trained for a greater number of epochs, as such, the model was trained for 1000 epochs. Due to the increased number of epochs, total training time lasted approximately twenty-four hours. Evaluation time remained consistent with RCAN and CARE models.

*Deep Learning Image Preprocessing*

Power- and gain-calibrated image stacks were loaded into Python as MAT data files. Image stacks featured an inconsistent number of depths (z-slices) due to patient-to-patient variability in cervical epithelial thickness. Due to differences in the number of z-slices, images were analyzed using a 2D network instead of a 3D network. Images with >10% of pixels considered to be cellular cytoplasm were selected for analysis and further normalized using percentile normalization as described previously[1,2]. Briefly, the images were rescaled such that the intensity values representing the lower two percent of the images were shifted to zero and the intensity values representing the upper 99.9 percent of the images were shifted to one. While the original implementation in *Weigert et al.* (2018) clipped values between 0 and 1, we did not clip negative pixel values and values greater than one[1].

Negative pixel values were clipped to zero by rectified linear unit layers in all models, while values greater than one can provide valuable high spatial frequency information and were preserved. Exclusion of the clipping step was in line with the implementation of percentile normalization in *Chen et al. (2021)*[2]. Stacks were normalized per slice and then split into four 256 x 256 images. The z-slices were appended to one another, generating, per image stack, a (4 x z-slices) x 256 x 256 image stack. For image reconstruction, additional information such as stack start and stack end locations were recorded per ROI. The stack start and end locations were later used to split training and validation sets based on ROI rather than randomly selecting different patches which represent a mixture of ROIs and depths. A training dataset and test dataset were generated independently with a consistent seed to ensure a consistent test dataset. For WU-net implementation, a 2D-DWT was applied to the patched 256 x 256 images using a biorthogonal 1.1 mother wavelet (greater detail can be found in Supplementary Discussion S2).

*Deep Learning Metrics*

Common metrics utilized in image restoration, super-resolution, and denoising were used to assess model performance before downstream analysis. PSNR and SSIM are validated metrics for such a task[1–3,7,10]. Briefly, PSNR is a metric that assesses the difference between the maximal true signal and the corruption of an image by noise[10]. As the difference between the corrupted image and true image is minimized, PSNR reaches higher values, indicating improved image quality[10]. PSNR was calculated using equation 1:

$$PSNR = 10 \cdot \log_{10}\left(\frac{MAX_I^2}{MSE}\right) \qquad (1)$$

Where $MAX_I$ is the maximum pixel intensity for an image (255 for an 8-bit greyscale image) and $MSE$ was calculated using equation 2:

$$MSE(x,y) = \frac{1}{m \cdot n} \sum_{i=0}^{m-1} \sum_{j=0}^{n-1} [x(i,j) - y(i,j)]^2 \quad (2)$$

Where $x$ is the ground truth image of size m x n and $y$ is the restored image of the same size. While PSNR is well characterized as a metric for image enhancement, it lacks assessment of perceived visual quality[3]. SSIM seeks to correct this by extracting the key features used in the human visual system to assess image quality. SSIM focuses on luminance, contrast, and structural differences between two images[3]. SSIM was calculated using equation 3:

$$SSIM(x,y) = \frac{(2\mu_x\mu_y + C_1)(2\sigma_{xy} + C_2)}{(\mu_x^2 + \mu_y^2 + C_1)(\sigma_x^2 + \sigma_y^2 + C_2)} \quad (3)$$

Where $x, y$ are the paired ground truth and restored image, $\mu_x$ and $\mu_y$ represent the mean intensity of the images, respectively, $\sigma_{xy}$ is the covariance of the two images, $\sigma_x^2$ and $\sigma_y^2$ are the variances of the two images' intensity, and $C_1$and $C_2$ are constants used to stabilize the division and are set based on a ratio of maximal image intensity and a stability constant[3]. SSIM values are calculated locally rather than globally using a circular-symmetric gaussian weighting function[3]. For all studies, a 3 x 3 filter with a sigma of 0.5 was used during loss calculation and an 11 x 11 filter with a sigma of 1.5 was used during metric calculation.
